# Injectable Janus Base Nanomatrix (JBNm) in Maintaining Long-Term Homeostasis of Regenerated Cartilage for Tissue Chip Applications

**DOI:** 10.1101/2024.10.05.616785

**Authors:** Anne Yau, Ian Sands, Wuxia Zhang, Yupeng Chen

## Abstract

Engineered cartilage tissues have wide applications in in vivo cartilage repair as well as in vitro models, such as cartilage-on-a-chip or cartilage tissue chips. Currently, most cartilage tissue engineering approaches focus on promoting chondrogenesis of stem cells to produce regenerated cartilage. However, this regenerated cartilage can dedifferentiate into fibrotic tissue or further differentiate into hypertrophic or calcified cartilage. One of the most challenging objectives in cartilage tissue engineering is to maintain long-term cartilage homeostasis. Since the microenvironment of engineered cartilage tissue is crucial for stem cell adhesion, proliferation, differentiation, and function, we aim to develop a novel scaffold that can maintain the long-term homeostasis of regenerated cartilage. Therefore, we developed a library of Janus base nanomatrices (JBNms), composed of DNA-inspired Janus nanotubes (JBNts) as well as cartilage extracellular matrix (ECM) proteins. The JBNms were developed to selectively promote chondro-lineage cell functions while inhibiting bone and endothelial cell growth. More importantly, the JBNm can effectively promote chondrogenesis while inhibiting hypertrophy, osteogenesis, angiogenesis, and dedifferentiation. Additionally, the JBNm is injectable, forming a solid scaffold suitable for producing and maintaining regenerated cartilage tissue in microfluidic chips, making it ideal for tissue chip applications. In this study, we successfully created cartilage tissue chips using JBNms. These chips can model cartilage tissue even after long-term culture and can also mimic arthritis progression, making them useful for drug screening. Thus, we have developed a novel nanomaterial approach for improved cartilage tissue engineering and cartilage tissue chip applications.

## Introduction

Cartilage tissue engineering holds great promise for the regeneration of lost or damaged cartilage because it leverages nanomaterial technology to redirect cell populations at the cellular level, forming functional tissue like natural occurring extracellular matrix (ECM)[1]. Cartilage ECM is a complex mixture of molecules that surround cells, tissues, and organs, consisting of different glycoproteins, proteoglycans, and other macromolecules including fibronectin (FN), matrix proteins (Matn), cartilage oligomeric matrix protein (COMP), laminin, and many other proteins. These components are responsible for orchestrating cell-cell interactions, mediating cell behavior, and providing structural and mechanical support to the tissue[2]. The ECM also serves as a reservoir for growth factors, cytokines, and various signaling molecules, contributing to its role in orchestrating cellular responses.

Strategies to regenerate damaged or diseased cartilage can be derived into two major focuses: (1) multipotent cells capable of proliferation and/or differentiation into cartilage and (2) a controlled and tunable microenvironment to influence cell behavior through physical and biochemical cues. The successful integration of stem cells and native tissue is contingent on our ability to mimic local physiological conditions and influence lineage commitment towards articular cartilage cells and ECM[3]. Previous tissue engineering endeavors for articular cartilage repair have focused on developing rigid synthetic and organic scaffolds that promote stem cell adhesion and attempt to integrate native and regenerated tissue. Unfortunately, stem cell lineage commitment is nuanced and becomes increasingly difficult when considering the delicate balance of chondrogenic, osteogenic, and hypertrophic signaling molecules regulated during tissue repair[4].

Modern biomaterials have evolved to show greater compatibility with cell and molecular biology, thereby providing biochemically relevant signals at targeted locations. They are often characterized by their ability to be integrated with cell and biomolecules to provide biochemical cues at a specific location. Biomaterials such as Matrigel and polymer-based nanomaterials have been leveraged to provide a suitable microenvironment for in vitro cartilage cell culture[5–7]. These biomaterial scaffolds furnish a three-dimensional (3D) framework that not only lends structural support, but also orchestrates the trajectory of cell and tissue growth, mirroring the functions performed by the native ECM[8].

One major limitation of current biomaterial scaffolds is the lack of homeostatic maintenance in regenerated cartilage[9]. Since many of the growth factors responsible for chondrogenic differentiation are also present in osteogenic differentiation pathways, regenerated cartilage tissue is subject to chronic-term de-differentiation or fibrosis[10–12]. This leads to molecular composition and mechanical property discrepancies between native and regenerated tissue which perpetuates cartilage degradation and can even promote local inflammation[13]. Therefore, it is critical that cartilage regeneration efforts focus on chondrogenic and hypertrophic gene expression to assess the long-term feasibility of biomaterial scaffolds for cartilage tissue regeneration. The ability to differentially regulate chondrogenic biomarkers while inhibiting the upregulation of hypertrophic proteins is essential for successful tissue repair.

Biomolecular or growth factor encapsulation is one method we can leverage in tissue regeneration scaffolds to improve healthy cartilage regeneration. Scaffolds can help direct stem cell proliferation and differentiation by incorporating growth factors or ECM-mimicking proteins to emulate natural molecular interactions. Although growth factor encapsulation is a viable method for chondrogenic differentiation, common growth factors like TGF-B will interact with chondrogenic and osteogenic differentiation pathways[14, 15]. This leaves the encapsulation of cartilage ECM mimics which include proteins like Matrillins, Fibronectin (FN), Collagens, and Laminins which can be incorporated into scaffolds to provide structural and functional maintenance to regenerated cartilage. Previous studies have demonstrated impact of ECM proteins not only for chondrogenic differentiation, but also for homeostatic maintenance to prevent the dedifferentiation of regenerated tissue[16]. This provides the incentive to develop novel biomaterial scaffolds that emphasize biocompatibility and biomolecular encapsulation to improve cell adhesion, proliferation, and the biosynthesis of crucial matrix molecules during tissue regeneration.

Herein, we develop a Janus Base Nanomatrix (JBNm); a biomimetic tissue regeneration matrix comprised of DNA-driven Janus nanotubes (JBNt) and various ECM proteins (such as matrilin and other cartilage-specific proteins). Unlike traditional synthetic scaffolds, JBNts are DNA-mimicking subunits synthesized through non-covalent interactions which enables exceptional biocompatibility. As a standalone scaffolds, JBNts exhibit a high density of outfacing lysine sidechains which facilitates molecular-level interactions with negatively charged cellular membranes. Apart from the prolific improvements to cell adhesion, the positively charged JBNts are an excellent template for negatively charged biomolecular encapsulation. JBNts are further functionalized with various cartilage specific proteins (Matrilins, COMPs, FN) through electrostatic forces to create a biomimetic matrix (JBNm) stabilized with structural ECM proteins for cartilage regeneration and maintenance. One additional advantage of JBNm lies in its mechanical properties as it self-assembles in aqueous solution. Unlike, rigid scaffolds, injectable scaffolds are able to adopt a pre-formed 3D porous structure before binding to biological agents and interfacing with the injured area through injection. As in vitro research advances towards complex three-dimensional cultures, models like tissue-on-a-chip are becoming more popular methods of therapeutic and diagnostic screening. Form-fitting regenerative matrices like JBNm hold a significant advantage over rigid scaffolds due to their injectability into hard-to-reach areas like damaged tissues and three-dimensional regeneration models.

The objective of this study is to develop a library of JBNms comprised of JBNts and different proteins found in the ECM, including Matrilin-1 (Matn1), Matrilin-3 (Matn3), COMP, and FN to promote chondrogenic lineage function in cartilage. We identified the most appropriate JBNm to promote the regeneration of cartilage tissue injected into tissue chips, thereby creating an ideal environment to support the chondrogenesis process of the chondro-lineage cells. We assessed the cell function of cartilage cells in tissue chips and demonstrated JBNms ability to promote stem cell chondrogenesis, maintain cartilage cell homeostasis, and prevent long-term hypertrophy, as well as prevent osteogenesis and other fibrotic differentiation lineages. The characterizations of the JBNm were observed via TEM, light microscopy, UV-Vis, and zeta potential. Morphology and effects of various JBNm in different cell types were studied: (i) the human mesenchymal stem cells (hMSCs), (ii) cartilage cells, (iii) osteoblast and (iv) human vascular endothelial cells (HUVEC). JBNm with Matn1 was selected for cartilage regeneration due to its highest anti-osteogenic and anti-angiogenesis characteristics resulting in one of the best JBNms for cartilage regeneration. The design aim of JBNm in cartilage tissue chips is to mimic 3D-like environment providing physiological microenvironment for cartilage cells to proliferate and generate ECM *in situ.* JBNm-in-tissue chip did not affect cell viability and did increase matrix synthesis, especially under dynamic loading where the upregulation of cartilage-specific genes like aggrecan (ACAN) Collagen 2 (COL2A1) and SRY box 9 (SOX9) were observed.

To demonstrate the potential applications of JBNm in therapeutic screening applications, we added Interleukin-1α (IL-1α) (5ng ml^-1^) on both JBNm and negative control (NC) samples to provide biochemical stimulation and analyze the release of glycosaminoglycans (GAG), followed by testing a series of anti-inflammatory drugs, Interleukin 1 Receptor Antagonist (IL-1 Ra) and Resveratrol with RT-qPCR. Lastly, the tissues’ ability to maintain stable internal conditions over extended periods of time, often through more complex regulatory mechanisms than short-term homeostatic responses were studied. hMSCs and chondrocytes were cultured over a period of 28 days and gene expression changes such as ACAN, COL2A1 and SOX9 were analyzed to determine the changes overtime. The success of JBNm-in-tissue chip will help pioneer future research including the incorporation of different proteins with various JBNts developed in our lab for therapeutic applications.

## Results

### 2.1 JBNt material characterization (**Figure 2**)

Janus base nanotubes (JBNts) are a family of Janus base nanomaterials (JBNs) self-assembled from G^C or A^T units mimicking DNA base pairs. The G^C monomer motif possesses Watson-Crick hydrogen bond ‘donor-donor-acceptor’ and ‘acceptor-acceptor-donor’ motifs present in guanine and cytosine, respectively (**Figure 2i**). The G^C monomers may also be conjugated with amino acid moieties that influence supramolecular chirality and electrostatic properties upon self-assembly (**Figure 2ii**). Supramolecular self-assembly occurs in aqueous solution as a result of G^C monomer interactions, creating a 6-member sp2 hybridized aromatic-ring system (rosette) with outfacing amino acid conjugates (**Figure 2iii**). Non-covalent interactions including hydrogen bonding, the hydrophobic effect, and the π-π stacking effect drives rosettes on top of one another forming a nanotubular structure with a high density of outfacing amino acids. These nanotubes (JBNts) are <1nm in diameter and can self-assemble upwards to multiple microns in length. Apart from the hydrogen bonding occurring between G^C monomers, H-bonds between stacked amino acid residues creates an inter-rosette hydrogen bond network responsible for stabilizing and preserving nanotube helicity. Another key feature of JBNt self-assembly lies between the aromatic rings that constitute the nanotubes core. JBNt monomers possess a two-ring aromatic core with each carbon possessing a pair of p-orbitals above and below its plane. Core aromaticity allows π electron mobility around the 10-carbon system, effectively creating π electron clouds below and above each assembled JBNt rosette. These π electron clouds drive the “base stacking effect” which is a phenomenon most commonly observed in the chemical structure of DNA base pairs that form the traditional double helix structure.

**Figure 1.**
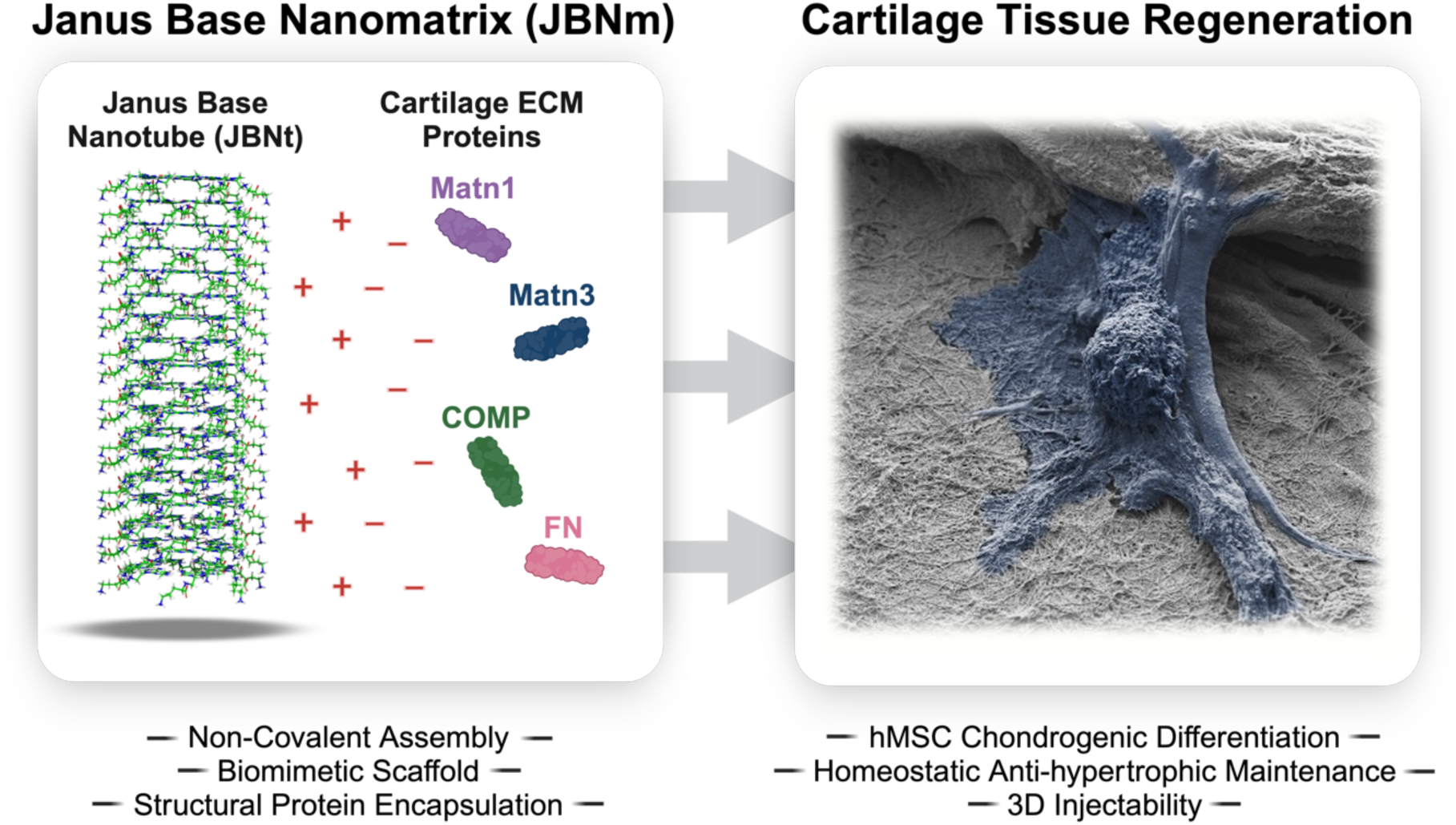
JBNts can be assembled in aqueous solution and mixed with cartilage matrix proteins to form a dense matrix for cell adhesion and differentiation (JBNm). JBNm provides an optimal platform for stem cell adhesion, chondrogenic differentiation, and homeostatic maintenance for healthy cartilage repair.

**Figure 2.**
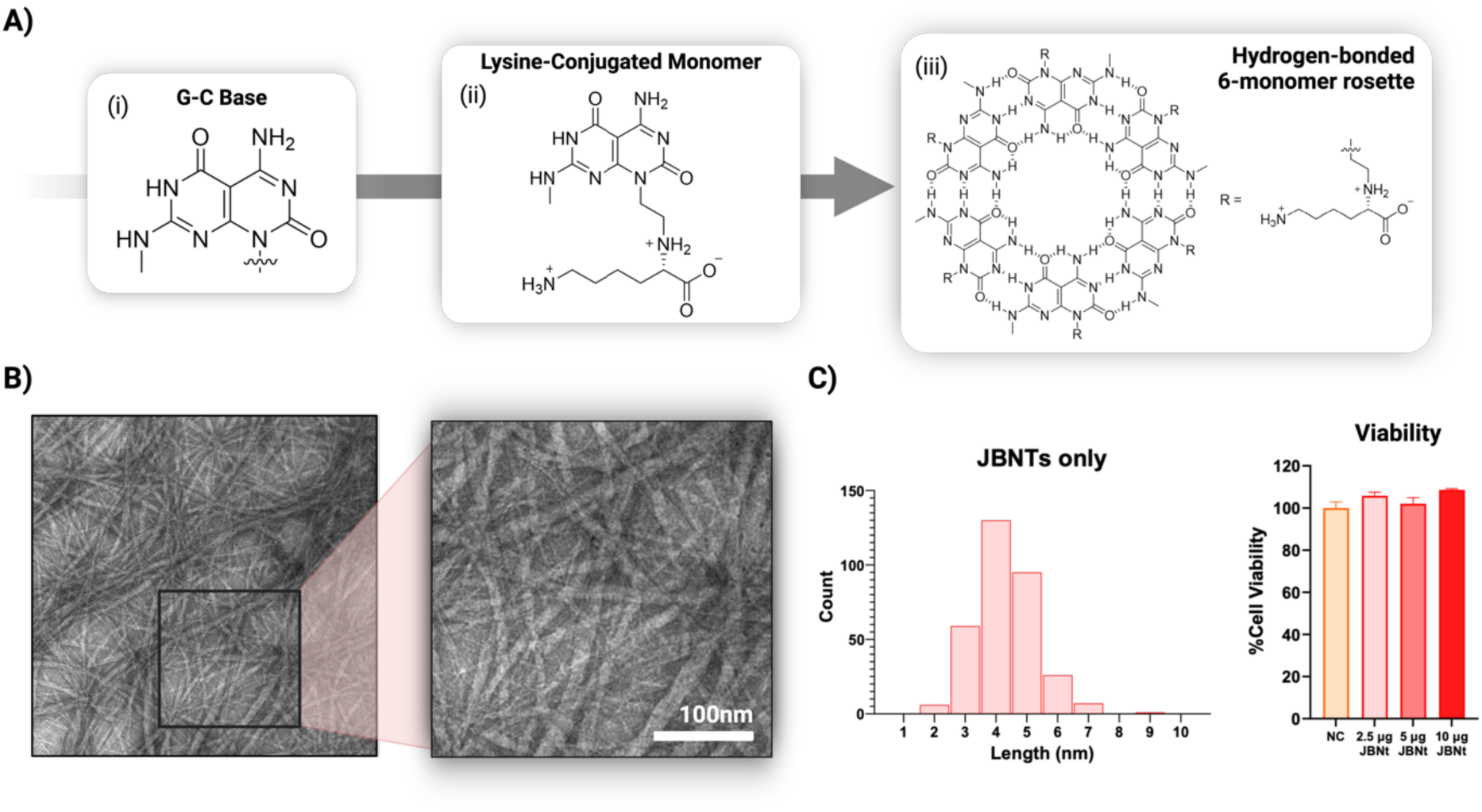
Material characterization of JBNt. **(A)** molecular structure of JBNts derived from DNA-mimicking monomers. **(B)** Transmission electron microscopy (TEM) image of JBNt. **(C)** Size distribution of the width of JBNt and the cartilage cell viability of JBNts up to 10 µg mL^-1^.

JBNts are observed in transmission electron microscope (TEM) and showed an average width of 4.3 ± 1.1 nm once diluted into single-strand solutions (**Figure 2B**) (**Supplemental Figure 1**). JBNts are positively charged and as expected, the zeta-potential of the 23 ± 2 mV at a concentration of 20 µg mL^-1^. JBNts show absorbance at 220 nm and 280 nm from the lysine side chains and aromatic rings of the Janus bases respectively which mimics the native collagen typically found in the ECM. We evaluated the cytotoxicity of JBNts seen in **Figure 2C**, culturing human cartilage cells with JBNts for 24 hours with concentrations of JBNts set as a gradient at 10 µg mL^−1^, 5 µg mL^−1^, 2.5 µg mL^−1^, and 0 µg mL^−1^ as NC. Even at 10 µg mL^−1^, the JBNts consistently presented excellent cell viability (>100%) when compared to the NC. The results confirmed that the DNA-inspired JBNts contributed to excellent biocompatibility.

### 2.2 JBNm and scaffold development for in vitro characterization

#### 2.2.1 Characterization of JBNms

The structure of the JBNm closely resembles fibrous structure of ECM in the human body and can act as collagenous material to provide support for cell adhesion. In this work, we selected four types of ECM proteins, Matrilin-1 (Matn1) in purple, Matrilin-3 (Matn3) in blue, Cartilage Oligomeric Protein (COMP) in green and Fibronectin (FN) in pink, to bundle with JBNts, creating JBNm for cartilage tissue formation in tissue chips. **Figure 3i-iii** shows the characterization of various JBNm in different ratios, starting with UV-Vis spectrophotometry, followed by the zeta potential and representative images of JBNm scaffolds seen in transmission electron microscope (TEM). In addition, **Figure 3iv** also shows the width distribution of each JBNm strand, increasing in size proportional to the protein concentration added. We first analyzed the JBNm assembly via UV-Vis and observed the bundling of the JBNts scaffold and proteins. The gradual decrease of absorbance peaks of the JBNm influenced by increasing concentration of protein demonstrated the formation of the JBNm between JBNts and proteins. When the JBNts were added with proteins of different concentration, the JBNt peaks decreased at 220 nm and 280 nm, indicating the bundling of proteins and JBNts, located in between each JBNts and protein spectra. The results demonstrate the coexistence of the bonding between JBNts and the proteins. The decrease of UV-Vis and zeta potential of JBNms when compared to JBNts supports the assembly of JBNm via non-covalent bond between JBNts and proteins.

**Figure 3.**
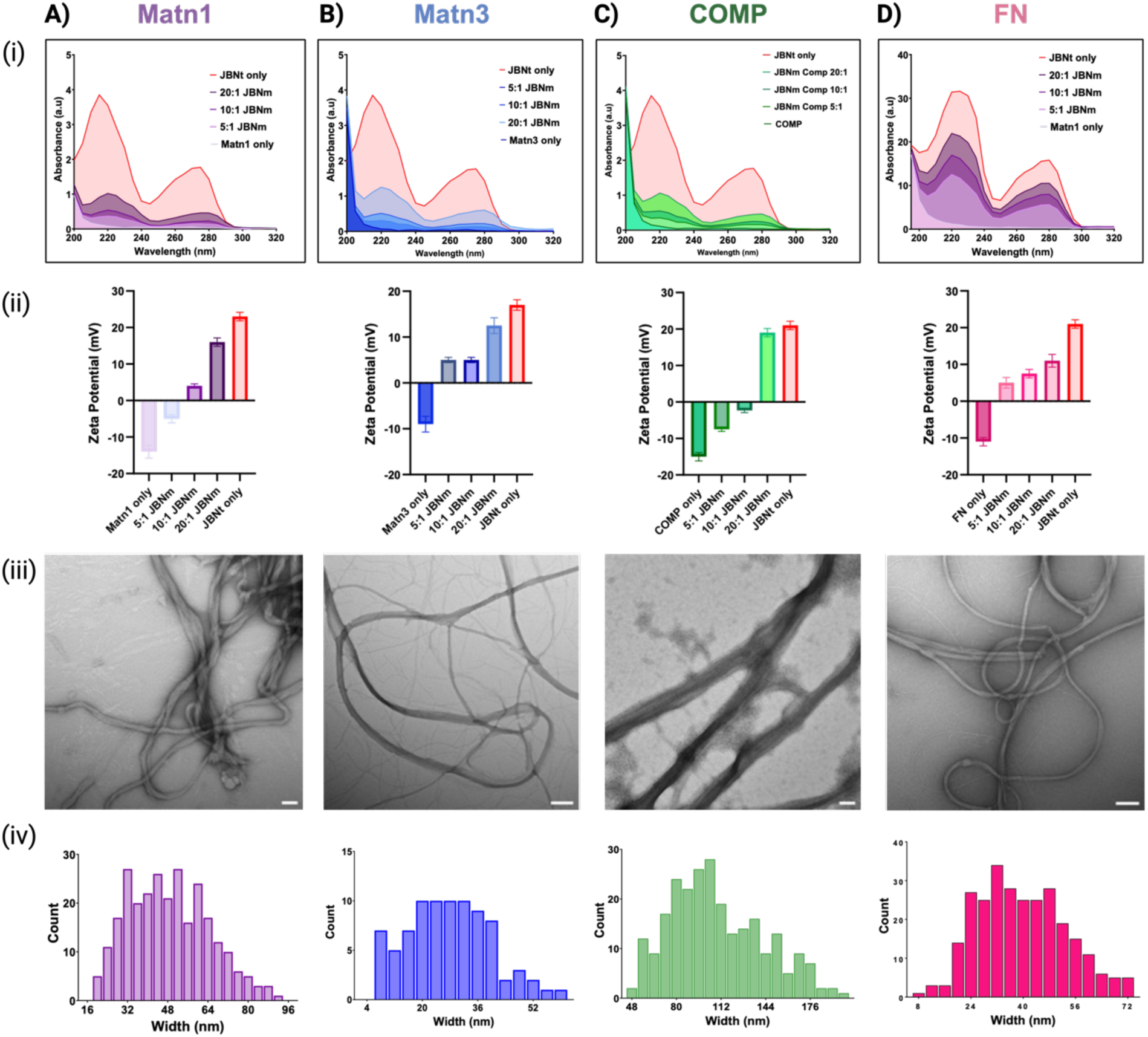
Characterization of JBNms. **(A)** Matn1 JBNm, **(B)** Matn3 JBNm, **(C)** COMP JBNm and **(D)** FN JBNm. **(i)** UV Vis of different JBNm comparing to JBNt (pink) at different ratio (5:1, 10:1 and 20:1) in terms of concentration in cell culture, **(ii)** Zeta potential or surface charge of JBNm nanomaterials in comparison to JBNt (pink), **(iii)** representative of TEM images of 10:1 JBNms and (iv) size distribution of the width of 10:1 JBNm.

ECM proteins are negatively charged based on their isoelectric point under physiological condition, as shown in **Figure 3ii**. In general, the JBNts are highly positive while the protein is negative in their respective charge[17]. The strong positive charge of JBNts, and the negatively charged of the proteins lead to the assembly of JBNts and proteins. The zeta potential and JBNm formation through images captured TEM (**Figure 3iii**). **Table 1** shows the zeta potential and size distribution of nanotube width comparison of each JBNm, JBNt and proteins.

**Table 1.** Catalog of JBNm synthesized with varying ratios of JBNt/ECM protein. Their ratios, zeta potential, and JBN width was captured using ImageJ.

| Matrix Composition | Ratio | Zeta Potential (mV) | JBN Width (nm) |
| --- | --- | --- | --- |
| JBNt only | - | 23 ± 2 | 3.9 ± 1.0 |
| Matn1 only | - | -14 ± 3 | - |
| JBNm/Matn1 | 5:1 | -5 ± 2 | 29.1 ± 9.6 |
| JBNm/Matn1 | 10:1 | 4 ± 1 | 21.7 ± 10.5 |
| JBNm/Matn1 | 20:1 | 16 ± 2 | 7.9 ± 3.7 |
| Matn3 only | - | -9 ± 3 | - |
| JBNm/Matn3 | 5:1 | 5 ± 1 | - |
| JBNm/Matn3 | 10:1 | 5 ± 1 | 26.6 ± 12.7 |
| JBNm/Matn3 | 20:1 | 12.5 ± 3 | - |
| COMP only | - | -15 ± 2 | - |
| JBNm/COMP | 5:1 | -7.5 ± 1 | - |
| JBNm/COMP | 10:1 | -2.3 ± 1 | 108.9 ± 26.2 |
| JBNm/COMP | 20:1 | 19 ± 2 | - |
| FN only | - | -11 ± 2 | - |
| JBNm/FN | 5:1 | 5 ± 2 | - |
| JBNm/FN | 10:1 | 7.5 ± 2 | 38.2 ± 13.9 |
| JBNm/FN | 20:1 | 11 ± 3 | - |

JBNm was fabricated in H2O at room temperature when proteins were added into JBNts. In the **Video Supplementary**, we added Matn1 to JBNt capturing the white floccules as soon as the addition of aqueous JBNts into Matn1 solution, forming cartilage-specific JBNm. Unlike hydrogels, JBNm is an injectable solid scaffold that provides microenvironments for many types of cells to adhere and grow. Their selectivity can be attributed to the specific ECM proteins that are likely to interact directly to cell membranes. These nanomaterials are not only selective depending on the cell type, but they also provide support for direct cell and tissue growth, mirroring the functions performed by the native ECM.

#### 2.2.2 JBNm improves MSCs and cartilage cell functions after 14 and 7 days

In this study, we cultured both human mesenchymal stem cells (hMSCs) and cartilage cells with different JBNm for 14 and 7 days, respectively, to determine the most appropriate JBNm for cartilage tissue formation. For hMSC culture, the positive control group was supplied with fresh TGF-β3 during chondrogenic medium change. The negative control group was not supplied with fresh TGF-β3, and was instead provided basal growth media. For cartilage cell culture, the negative control has no JBNm. In both cell-type cultures, chondrogenic and homeostasis markers such as COL2A1, ACAN, and SOX9 were measured and analyzed. We also looked at hypertrophic markers like Collagen Type X (COL10A1) and Indian Hedgehog (IHH) as well as Collagen Type I (COL1A1) after 7 and 14 days.

The selectivity of different JBNm formulations to multiple cell types was observed through morphological changes. **Figure 4A** shows that different JBNms influenced different cell-types compared to JBNts alone and negative control without any additives. For example, the JBNt/Matn1 JBNm promoted hMSC and cartilage cell functions while inhibiting osteoblasts and HUVECs, followed by the selectivity of JBNt/Matn3 JBNm, which shows similar trends. Matrilines regulate interactions between collagen fibrils and other matrix components, such as aggrecan, COMP, and different collagen types. Both Matn1 and Matn3 are essential during chondrogenesis[18]. COMP, on the other hand, is a type of glycoprotein that interacts with the ECM in cartilage and other tissues that plays a critical role in the structural organization of the ECM where it modulates Collagen 1 synthesis[19], in which we observed an upregulation of Collagen 1 seen in **Figure 4B-C**. On the other hand, FN is a glycoprotein used for cell attachment and binds through cell-surface receptors which influences cell shape, phosphorylation of intracellular proteins, and gene expression and is important in chondrogenic/osteogenic differentiation[20]. JBNt/FN JBNm promoted hMSC and HUVEC functions but inhibited osteoblasts and chondrocytes, which may be used for angiogenesis and soft tissue repairs.

**Figure 4.**
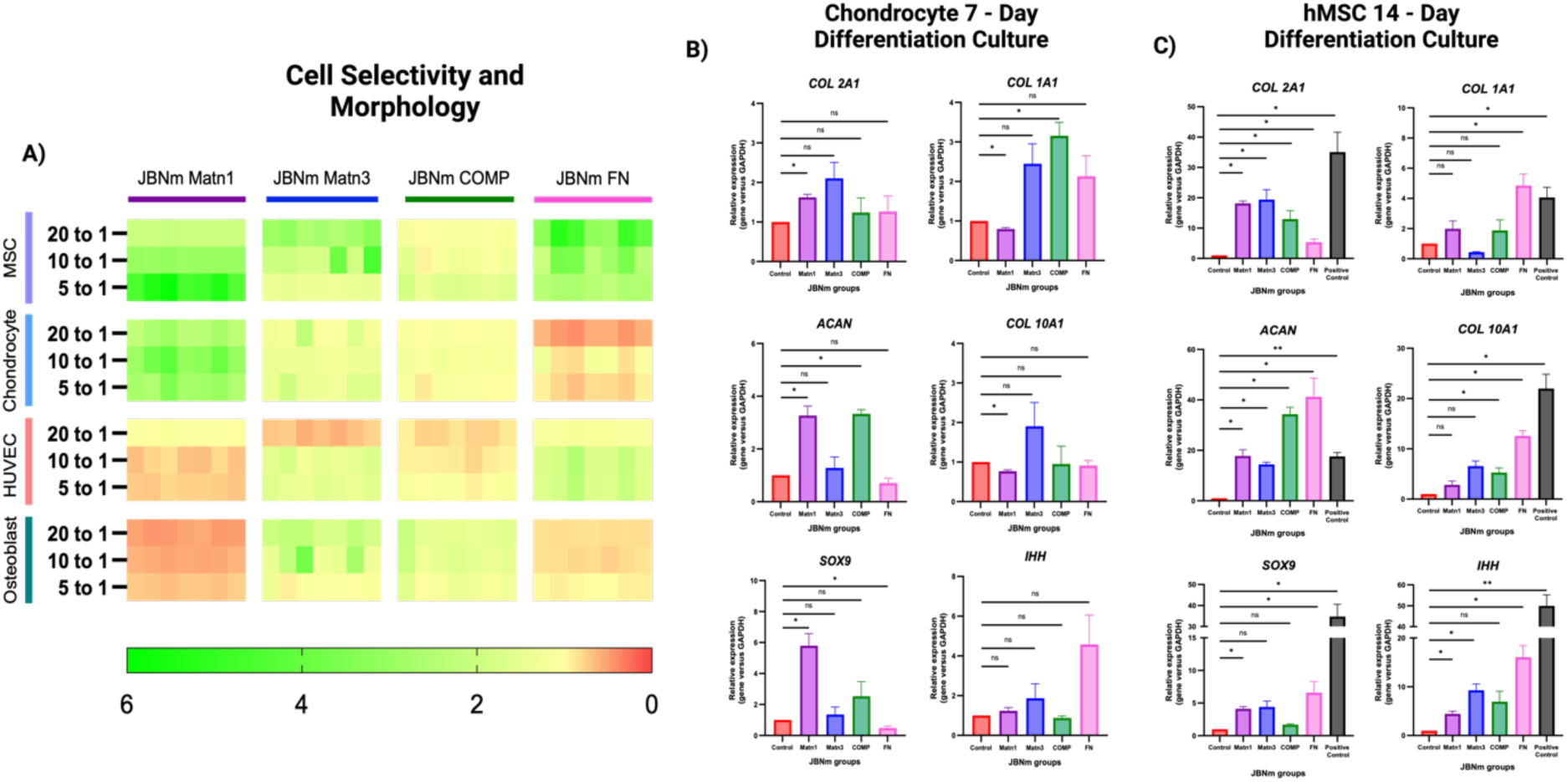
**(A)** Cell morphology properties such as compactness, eccentricity, aspect ratio, perimeter and maximum radius are obtained and analyzed with Cell Profiler, comparing different ratios of JBNm to negative control cultured on MSCs, chondrocytes, HUVEC and Osteoblast. All JBNm improved cell morphologies of MSC, however, only Matn1 JBNm and Matn3 JBNm improved cell morphology of cartilage cells (chondrocytes), while FN JBNm improved HUVEC. **(B-C)** PCR data of cartilage cells and hMSCs after culturing for 7 days, respectively, in engineered tissue constructs. Both results showed Matn1 and Matn3 JBNm encouraged chondrogenesis with upregulation of COL2A1, ACAN and SOX9.

After culture, total RNA was extracted from hMSCs, and cartilage cells and rt-qPCR was carried out for gene analysis of the functional outcomes of the JBNms. At first glance, the positive control was highly upregulated in all expressions, including COL2A1, ACAN, SOX9, COL1A1, COL10A1 and IHH, which is as expected. COL2A1 is a major component of the cartilage ECM and expressed in chondrocytes, providing cartilage with its tensile strength while ACAN, a large proteoglycan molecule, helps maintain cartilage’s compressive resistance. Both COL2A1 and ACAN are regulated by SOX9. In this study, JBNm Matn1 and JBNm Matn3 groups increase COL 2A1, ACAN and SOX9 expressions, representing the chondrogenesis markers while COL1A1, COL10A1, and IHH expression were not significantly upregulated, representing the hypertrophy markers, in both cell cultures. More importantly, these groups presented better bioactivity than the positive control group indicating chondrogenesis is prevalent in JBNm groups.

In a 14-day hMSC culture with differentiation medium, all gene expression of the positive control were upregulated, including the chondrogenesis and hypertrophy markers, which were expected. However, hypertrophic markers such as the COL10A1 and IHH, as well as fibrosis marker (COL1A1) expression were lower when cultured in JBNms, except COL10A1 expression in JBNm FN. COL2A1 are significantly upregulated in all JBNm, which is expected because proteins in JBNms are all cartilage ECM protein. When cultured with cartilage cells, only the JBNm Matn1 group significantly upregulates the expression of COL2A1 compared to negative control. Again, these results were as expected since Matn1 is a protein found prominently in cartilaginous tissues, and its role in maintaining structural integrity of cartilage. A similar trend of ACAN was observed when cells are cultured in hMSCs (**Figure 4C**) and on cartilage cells (**Figure 4B**). Interestingly, although SOX9 was upregulated in hMSCs culture with JBNm Matn3, they are not significantly upregulated in cartilage cells. These results confirm that JBNm Matn1 and JBNm Matn3 are most suitable JBNm for tissue regeneration for our next experiments.

### 2.3 JBNm aids in facilitating cells adhesion and proliferation

The bioactivity of the JBNm in cartilage cells was further analyzed to determine its feasibility in tissue chips using JBNm Matn1. **Figure 5** shows the quantitative morphology of chondrocyte cultured after 12 hours in chamber glass coated with different materials, such as non-additive negative control (NC), JBNt, JBNm and Matn1. We quantified the cell morphology parameters through Cell Profiler and conducted statistical analyses to assess the effects of each JBNm. The statistical analysis showed that JBNm groups have the highest bioactivity and have the greatest effect in all groups, including cell surface and angle ratio (**Figure 5b**).

**Figure 5.**
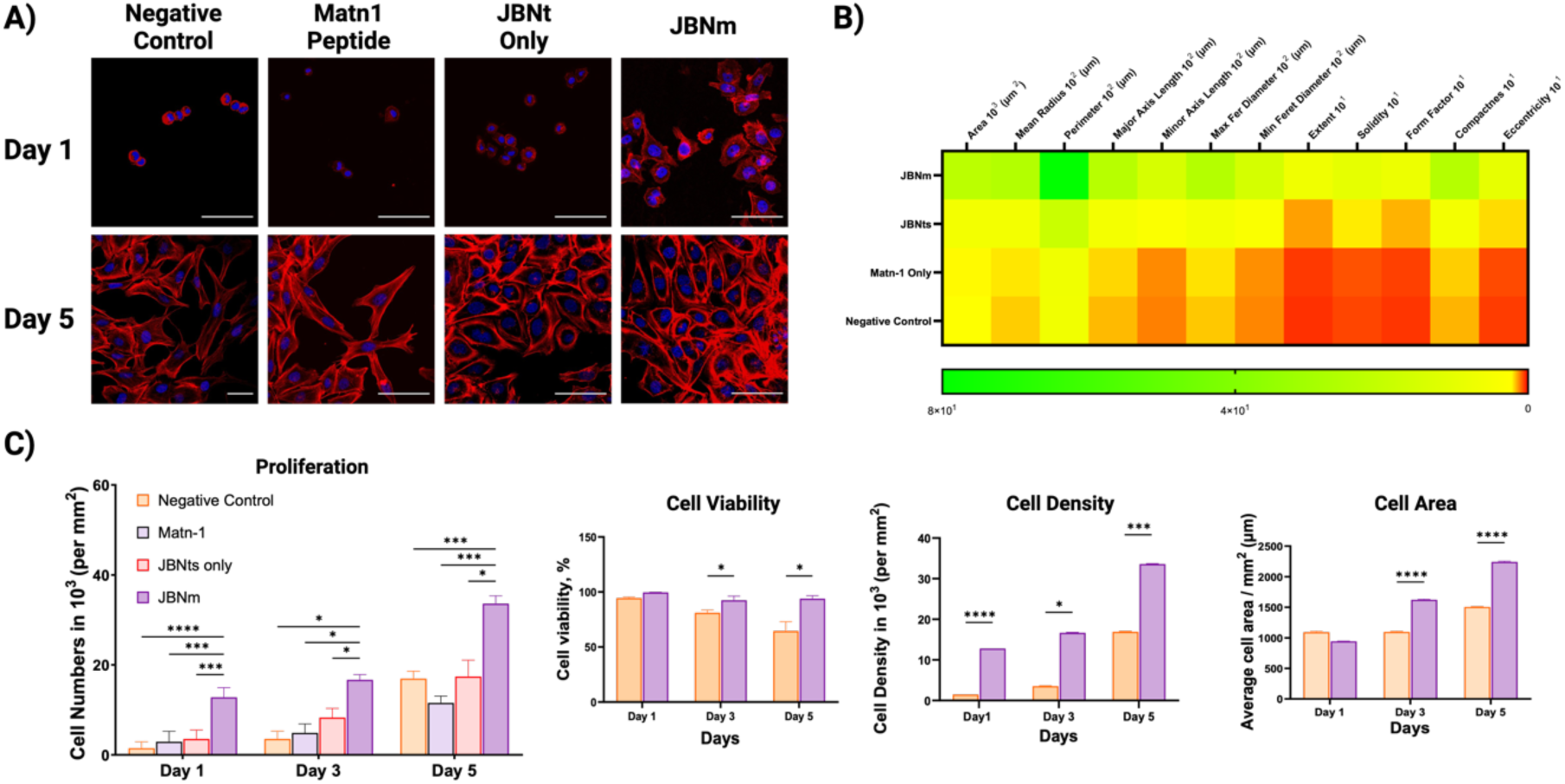
Confocal images, quantitative and statistical analysis of cell morphology and cell proliferation. **(A) (i)**. Negative control only without any additives; **(ii)**. Matn-1 only; **(iii)**. JBNts only; **(iv)**. JBNm at 10:1 ratio. **(B)** Statistical analysis map of cell shape parameters for C28/I2 between different groups and compared with negative control (NC). These data are obtained from CellProfiler. N=30. *P ≤ 0.05, **P ≤ 0.001, ***P ≤ 0.0001, ****P ≤ 0.00001. N=3. Starting cell count 2000 cells. Scale bar = 100 µm. This shows the heat map of chondrocyte cell shape between different groups and compared to negative control without any additives. The cells were seeded onto a JBNm coated chamber slides overnight before fixing and staining and cell parameters such as the area, mean radius, perimeter, major axis length and others were significantly enhanced supporting the hypothesis that JBNm can improve chondrocyte bioactivity; **(C)** Cell proliferation of cartilage cells on tissue culture plate after incubated with different materials (10:1 JBNms, Matn-1 only, JBNts only and negative control without additives) for 1 day, 3 days and 5 days, respectively.

The qualitative morphology of the cell adhesion grown on different materials and different time points (Day 1 and 5) was observed (**Figure 5**), showing excellent binding affinity to JBNm. JBNm ratio of 10:1 showed higher cell adhesion compared to other groups such as NC, Matn1 only, and JBNt only. We also analyzed cell proliferation at different times. The effectiveness of monolayer cell culture on JBNm-coated chamber slides shows that cartilage cells prefer the attachment and proliferation on JBNm substrates as opposed to no additives. Although Matn1 is a cartilage ECM that would promote chondrogenesis, high concentration of Matn1 does not seem to increase the proliferation of cartilage cells as much as JBNt alone or JBNm samples (**Figure 5C**). The cellular proliferation can also be explained with the characterization of JBNm where the zeta potential of 10:1 JBNm is positive, while 5:1 is negative. Based on our previous experiments, substrates with negative zeta potential do not promote cellular functions due to repulsion resulting from negatively charged tissues or cells. With respect to other cellular response metrics (viability, density, membrane area), JBNm provided a significant improvement, with the most notable discrepancies at Day 5.

### 2.4 Injectable JBNm scaffolds promote cartilage tissue development in tissue chip mimicking healthy native cartilage tissues

#### 2.4.1 Validations of tissue chip

To more accurately mimic in vivo physiological conditions and demonstrate the advantages of JBNm as an injectable matrix, a three-dimensional cartilage-on-a-chip model was developed for in vitro testing. The fabrication and the setup of the tissue chip can be seen in **Supplementary Figure 2**. The fabrication of negative master mold utilizes the rapid prototype which is low in cost, quick and easy way in less than an hour. A negative master mold design obtained from SolidWorks. The device incorporated three channels, that are interconnected allowing for nutrients exchange among all channels. After the injection of cells and JBNm embedded in 1% agarose into the middle channel, the culture medium was perfused through both side microchannels at perfusion rate of 250 µl hr−1 to mimic fluid flow of nutrients in the human cartilage (**Figure 6A**). PDMS was used to make the chip for its optical properties and its bioinert nature, so any experiment could be clearly viewed under a microscope with no adverse side effects from interaction with the material [21]. The device incorporated three channels, that are interconnected allowing for nutrients exchange among all channels. After the injection of JBNm into the middle channel, the culture medium was perfused through both side microchannels at perfusion rate of 250 µl hr−1 to mimic fluid flow of nutrients in the human cartilage. PDMS was used to make the chip for its optical properties and its bioinert nature, so any experiment could be clearly viewed under a microscope with no adverse side effects from interaction with the material [21].

**Figure 6.**
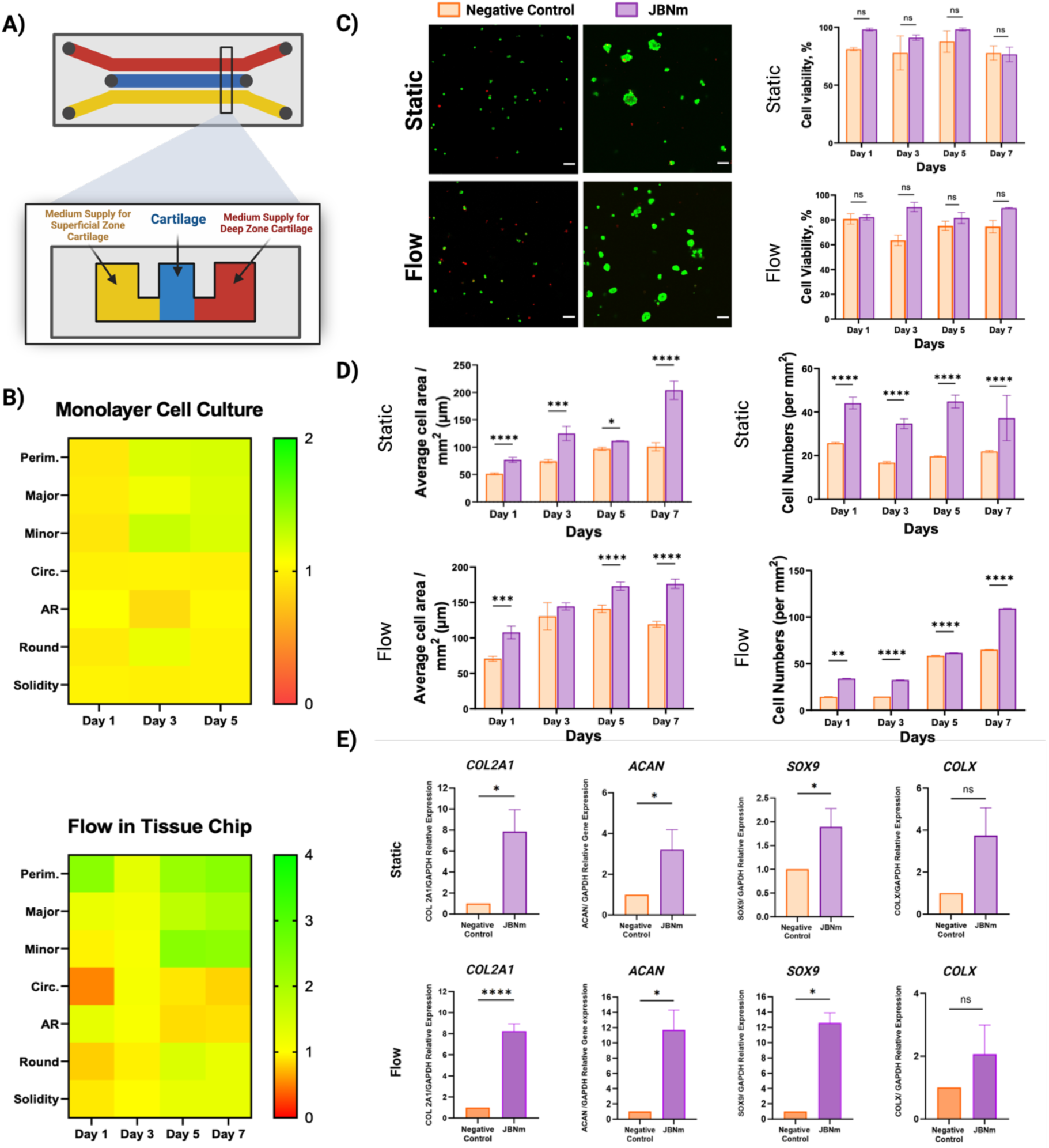
**(A)** Confocal images of live (green) and dead (red) cells comparing Negative control with no additives and JBNm as well as static to dynamic environment at 250 µL hr-1. **(B)** Statistical analysis of cell viability (i-ii), cell adhesion (iii-iv) and cell proliferation of monolayer cell culture (v-vi). The cell culture in tissue chip in static and dynamic environment at a flow rate of 250 µl hr-1. Calculation in this image is obtained from software ImageJ, (i-iii) cell viability, average cell are and total cell numbers, respectively. These data are obtained from CellProfiler. N=30. *P ≤ 0.05, **P ≤ 0.001, ***P ≤ 0.0001, ****P ≤ 0.00001. N=3. Scale bar = 100 µm. **(C)** Comparison of JBNm over negative control with no additives on all parameters such as perimeter, major and minor which signifies adhesion, circularity, aspect ratio, roundness and solidity which are also indicators for cell morphology. **(D)** RT-qPCR samples of JBNm and NC under static (i-iv) and flow condition (v-viii), “showing” that JBNm increase chondrogenesis markers such as COL2A1, ACAN and SOX9 while “did not upregulate COLX” markers.

We investigated cell morphology in the tissue chip design compared to a standard monolayer culture. The perfused tissue chip model improved multiple cell morphology metrics associated with adhesion, migration, and spreading (**Figure 6B**). We also explored cell viability in the tissue chip as well as the ability of JBNm to improve cell morphology and cell proliferation comparing the static (no flow) and a dynamic (with flow at 250 µL hr−1) environment. There was no significant difference on the viability of the cells when JBNm is injected into the chip and cultured in a static condition, compared the cells seeded in a condition without JBNm (**Figure 6C**). The cell viability remained similar when compared to negative control with no significant differences, although JBNm groups presented higher live cell counts compared to negative control with no additives.

The JBNm group has consistently shown significant effects on cell area and number regardless of the external stimuli (**Figure 6D**). This suggests that JBNm has a positive effect on the bioactivity of cells, leading to increased cell proliferation and differentiation. The area and number of cells often indicate the adhesion and proliferation activity of cells in the cultured state [22]. Interestingly, cell growth area on a monolayer culture was observed to be larger than the cells cultured in tissue chips. We speculate that the increase in growth area is due to both the adhesion to the 2D coated JBNm and the lack of vertical space when compared to a 3D microenvironment in a tissue chip [23]. Cell cultures on chips enable cells to expand in three-dimensional rather than two-dimensional cell culture on tissue culture plates (TCPs).

Due to shear stress from the fluid flow on both sides of the cartilage tissue, the cell sizes in our flow environment are lower than of static environment. This supported our viability study where the viability in the static condition is slightly higher than the cells subjected to dynamic loading. Although the cell viability between JBNm and NC was not significantly different in the dynamic chip with a flow rate of 250 µl hr-1, the viability cell percentage in the condition was lower when compared to static condition. This is expected due to the shear stress affecting the cells, causing slightly more cell deaths in dynamic condition [24].

#### 2.4.2 JBNm encourages chondrogenesis and cartilage tissue modelling

The aim of the tissue chip design was to establish a smaller scale model of healthy human cartilage tissues. In our experiment, we observed that cartilage cells tend to accumulate in the JBNm group due to the favorability of the cells towards the scaffold when compared to negative control (**Supplementary Figure 3**) which can also be seen in a study by Libo et. al. [25]. As expected, although cell adhesion is lower in dynamic condition than that of static condition, the cell numbers were higher in the dynamic condition indicating that cartilage cells prefer dynamic loading environment to function. The cell morphologies for tissue constructs in tissue chips were calculated using the open-source program, ImageJ, to examine the adhesion and proliferation of cartilage cells cultured with and without dynamic loading at a rate of 250 µl hr−1 observed in Figure 6C. Control devices were cultured without JBNm under flow of 250 µL hr-1. The JBNm group exhibited a higher number of cells due to the availability of JBNm as scaffold for cells to adhere to, supporting their proliferation activity. **Figure 6B** shows that JBNm showed improvements on all parameters such as perimeter, circularity, aspect ratio, roundness and solidity which signifies adhesion and an indicator for cell morphology. We observed a 2x fold change between JBNm and negative control in a monolayer cell culture in terms of cell morphology, however, a four-fold increase when they are cultured in a 3D environment. Cell morphology parameters such as perimeter, major and minor axis of cells were obtained analyzed in Cell Profiler and compared between JBNm and negative control.

Matn1 has been found to play a role in chondrogenesis, the process of cartilage formation including articular cartilage and growth plate cartilage. It plays a crucial role in maintaining the structural integrity of cartilage by interacting with other ECM components such as collagen and proteoglycans. It can influence chondrocyte differentiation and function, making it a potential candidate for promoting chondrogenic differentiation of stem cells when designing tissue-engineered cartilage constructs. Similarly, cartilage tissue constructs with injected JBNm were assessed with real-time qPCR, to test the functional outcomes of JBNms in tissue chips after 7 days in dynamic loading. As expected, chondrogenic markers, such as COL2A1, ACAN and SOX9 were upregulated seen in **Figure 6E**. We further characterized the effect of JBNm in tissue chips with and without dynamic loading, comparing them with and without JBNm on the production of GAG. The tissues were harvested to determine the total DNA and GAG in the tissue. The GAG and DNA content of the cartilage cells were determined using the DMMB assay and Hoechst 33528 assay analyzed in microplate reader at absorbance of 525 nm and with fluorescence with excitation and emission at 360-430 nm, respectively). GAG and DNA content of cartilage cells cultured in the tissue chip in dynamic loading showed significant improvements of GAG production in the JBNm injected chip compared to the negative control after 7 days (**Supplemental Figure 4**). In both NC and JBNm tissue chips, the proliferative effect of cells in the static condition was expected to be less than that of cells in the flow condition. JBNm groups has shown higher proliferation activity and GAG production after 14 days in both static and dynamic condition. The expression of specific cartilage proteins and genes, such as ACAN, SOX9, and COL2A1 (**Figure 6E**), is upregulated higher than NC in JBNm samples after 7 days of culture, supporting an increase in GAG synthesis in the JBNm group observed in this study.

### 2.5 JBNm prevents degradation of cartilage matrix proteins in a diseased models demonstrated in a cartilage tissue chip

#### 2.5.1 JBNm prevents further degradation of cartilage tissue after the addition of interleukin-1α within cartilage tissue

Inflammatory cytokines have been reported to stimulate cartilage matrix erosion in vitro [24]. Here, the degradation of cartilage tissue was observed when one channel of the cartilage tissue chip was subjected to IL-1α treatment while the other was not. The channel subjected to IL-1α treatment represents the cartilage tissue in the superficial zone, whereas the channel without no IL-1α treatment represents the deep zone shown in the schematic diagram. Then, the cartilage cells in the devices are cultured for 24 hours before subjected to treatment of 5 ng mL−1 of IL-1α into one channel out of the two denoting the superficial zone channel, at the flow rate of 250 µl hr−1 for 48 hours. The supernatants were collected at specific timepoints and measured with microplate reader at absorbance of 525 nm. We observed that the negative control cartilage tissue chip without JBNm presented excessive GAG degradation which typically is seen in many common joint diseases such as rheumatoid arthritis (RA) and osteoarthritis (OA) [26].

### 2.5. Determine the potency of drugs in inhibiting cartilage degradation using JBNm in cartilage tissue chip

Our JBNm has been shown to successfully help prevent further degradation of GAG and its’ release into supernatant, especially after the addition of IL-1α drugs. In the drug screening process, a biomechanical-based model based on a dose of pro-inflammatory cytokines in addition to the anti-inflammatory drugs were applied into the superficial zone of the tissue chip model to determine the pharmacological effects of JBNm in cartilage tissue chip. The experiments included a preliminary drug screening of the effect of two selected drugs IL-Ra (5 ng mg-1, 20 ng mL-1, 40 ng mL-1, 100 ng mL-1 and 200 ng mL-1) and resveratrol (0.1 µg mL-1, 0.5 µg mL-1, 2µg mL-1, 10 µg mL-1 and 20 µg mL-1) containing JBNm in tissue chips. After the cells in the tissue chips mature for 24 hours (Day T-1), they are subjected to treatment of pro-inflammatory cytokine for another 24 hours (Day T0). Then the tissue chips were treated with different dosages of drugs in addition to continuous IL-1α for the next 72 hours after that (Day T1-3) (**Figure 7A**). NC with constant supply of the anti-inflammatory drugs and IL-1α were used as negative controls while tissue chips with JBNm only were used as positive control. Supernatants were collected at selected timepoint and analyzed at the end of treatments on Day 3. As time goes by, the GAG degradation increased, leading to high GAG degradation seen at the end of treatment. GAG degradation was the most prominent in the positive control where only IL-1α was added. The treatment of anti-inflammatory drugs, either IL-Ra or Resveratrol successfully suppressed the GAG degradation which was as expected. The JBNm sample also exhibited the ability to reduce GAG degradation, which means higher integrity of cartilage tissue was present in the tissue chip. The release of GAG was based on the pre-treatment of Day 1 used as a baseline. A daily GAG release over the course of three-day IL-1α and drug treatments was observed in **Figure 7Bi-ii and 7Ci-ii**. Both **Figure 7Bi-ii and 7Ci-ii** showed that the GAG degradation in all groups (n=3) except for the IL-1α group increased starting on Day 0 when the IL-1α was added. Degradation rates dropped on Day 4-5 denoting the drugs have successfully suppress the degradation. IC50 was calculated with graphing and statistical software Graphpad Prism 9.0 and found to be 13.15 ng mL-1 for IL-Ra drug and 2.63 ug mL-1 for Resveratrol. This shows that JBNm can help create the homeostatic and chondrogenic microenvironment which are preferable and mimic cartilage sample better than without JBNm.

**Figure 7.**
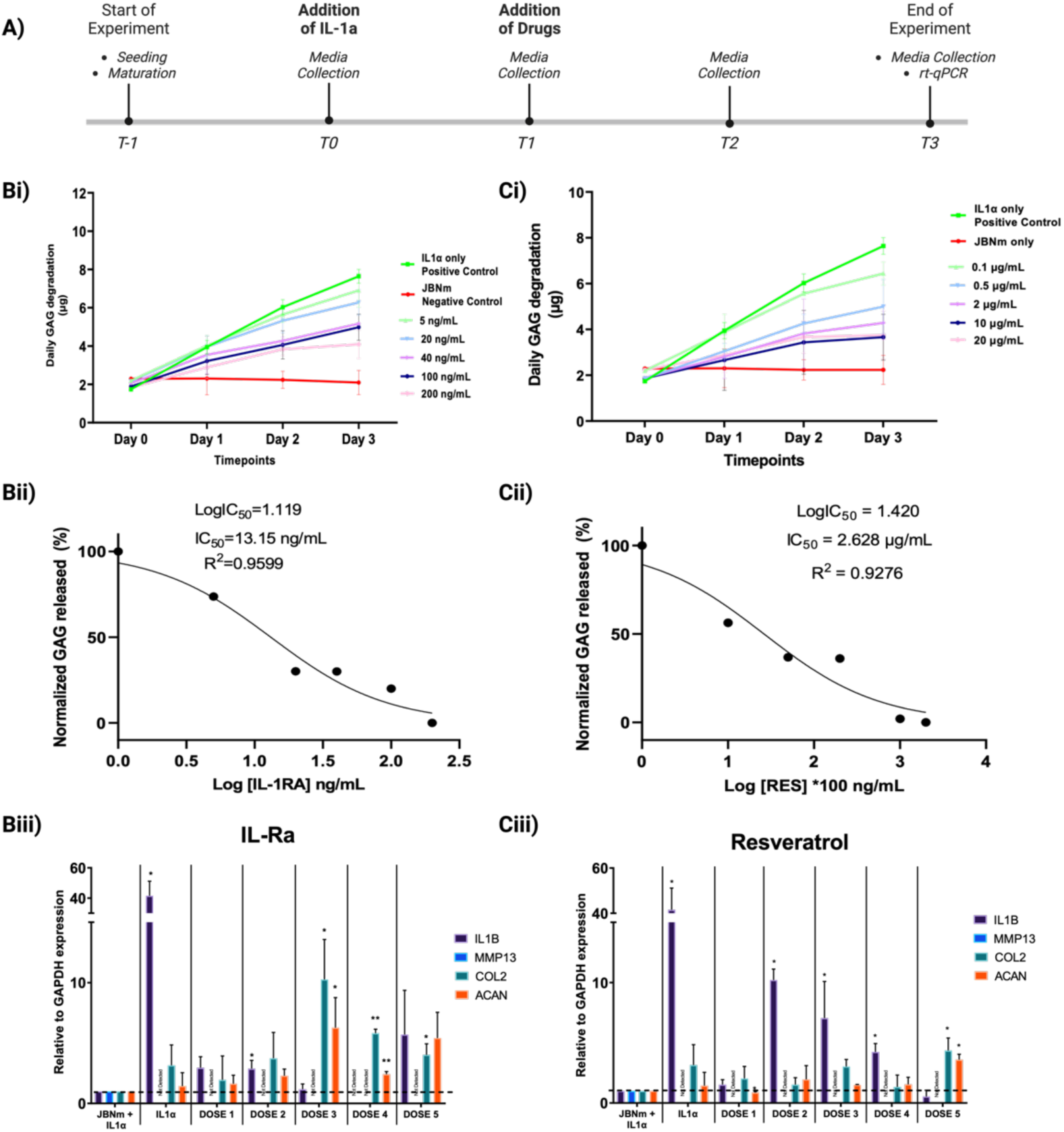
The experiments included a preliminary drug screening of the effect of two selected drugs IL-Ra (Dose 1 = 5 ng mg-1, Dose 2 = 20 ng mL-1, Dose 3 = 40 ng mL-1, Dose 4 = 100 ng mL-1 and Dose 5 = 200 ng mL-1) and resveratrol (0.1 µg mL-1, 0.5 µg mL-1, 2µg mL-1, 10 µg mL-1 and 20 µg mL-1) containing JBNm in tissue chips. **(A)** Shows the timeline of the study. After the cells in the tissue chips mature for 24 hours (Day T-1), they are subjected to treatment of pro-inflammatory cytokine for another 24 hours (Day T0). Then the tissue chips were treated with different dosages of drugs in addition to continuous IL-1α for the next 72 hours after that (Day T1-3). **(B-C)** IL-Ra and Resveratrol drugs on GAG release after every 24 hours **(i)**, IC50 calculations on both IL-Ra and Resveratrol **(ii)**, and RT-qPCR study on cartilage tissue after different dosage treatments, with NC with IL-1α sample as negative control.

In a diseased condition, cells secrete various inflammatory growth factors such as the TNFα and IL-1β into the surrounding area [27]. In pathophysiology of osteoarthritis pathway, the secretion of IL-1β and TNFα occurs and will lead to the degradation of cartilage caused by the upregulation of MMPs, and down regulation of ACAN and COL2A1. Therefore, we “mimic” the diseased condition by adding IL-1 α (5ng mL-1) and tested for matrix degradation in a biochemical assay, gene expression of matrix degradation (ADAMTS5 and IL-1β) and gene expression of chondrogenesis (ACAN and COL2A1) during the 3-day treatment. High GAG content indicates higher matrix degradation in tissue samples. From the data obtained, IC50 of the drugs for the study can be obtained as well (Figure 7 C-D). The cells were then subjected to 24 hours (Day 0) of IL-1α before treatment of different doses of drugs for 72 hours (Day 1-3). As seen in **Figure 7Biii**, the control with just IL-1α addition has high IL-1β gene expression. As the dosages for both drug treatments increased the gene expression of IL-1β decreased. However, the expression of IL-1β of dose 5 (200 ng mL-1) of the IL-Ra drug increased slightly which can be attributed to cytotoxicity at that dose concentration.

At the end of drug screening process, the samples were tested for gene expression study with RT-qPCR and JBNm with IL-1α sample as negative control. The samples were used to test for inflammatory gene expression, interleukin-1β (IL-1β), breakdown of ECM, a disintegrin and metalloproteinase with thrombospondin motifs 5 (ADAMTS5), and chondrogenic genes expression, COL2A1 and ACAN. **Figure 7Biii and 7Ciii** shows how the drugs affect the samples after 3 days of treatment, with different doses. The control used in the experiments was JBNm sample subjected to IL-1α treatment without any drugs to aid in suppressing the breakdown of GAG. The trend of COL2A1 and ACAN starting from dose 2 of IL-Ra, increased as the doses of IL-Ra increased can be observed until dose 5, where there is a slight down-regulation of COL2A1 and ACAN possibly due to toxicity. There is an up-regulation of IL-1β was observed in the IL-Ra treatment, indicating the possibility of high dosage IL-Ra at 200 ng mL-1. A similar trend was observed with Resveratrol treatment where there is an observable down regulation of IL-1β and upregulation of COL2A1 and ACAN as doses increased. The expression of ADAMTS5 of the control is normalized to one and there is an up-regulation of ADAMTS5 and IL-1β starting on dose 1 with continued downregulation as doses increases.

#### 2.5.3 JBNm cartilage tissue chips for long-term tissue-chip study

Among the various hurdles associated with cartilage tissue chips is the limitation that when subjected to biochemical or mechanical stimulation, tissue samples within these chips cannot be reused and struggle to accurately mimic the physiological and pathological conditions of native cartilage tissue. The potential for long-term cultures opens up the opportunity for repetitive exposure to substances, addressing this limitation. In this study, we have performed a 28-day culture study using cartilage cells and hMSCs. Seen in **Figure 8A and 8B**, gene expressions of cartilage homeostasis such as COL2A1, ACAN and SOX9 as well as gene expression for hypertrophy such as IHH, COL10A1 and dedifferentiation like COL1A1.

**Figure 8.**
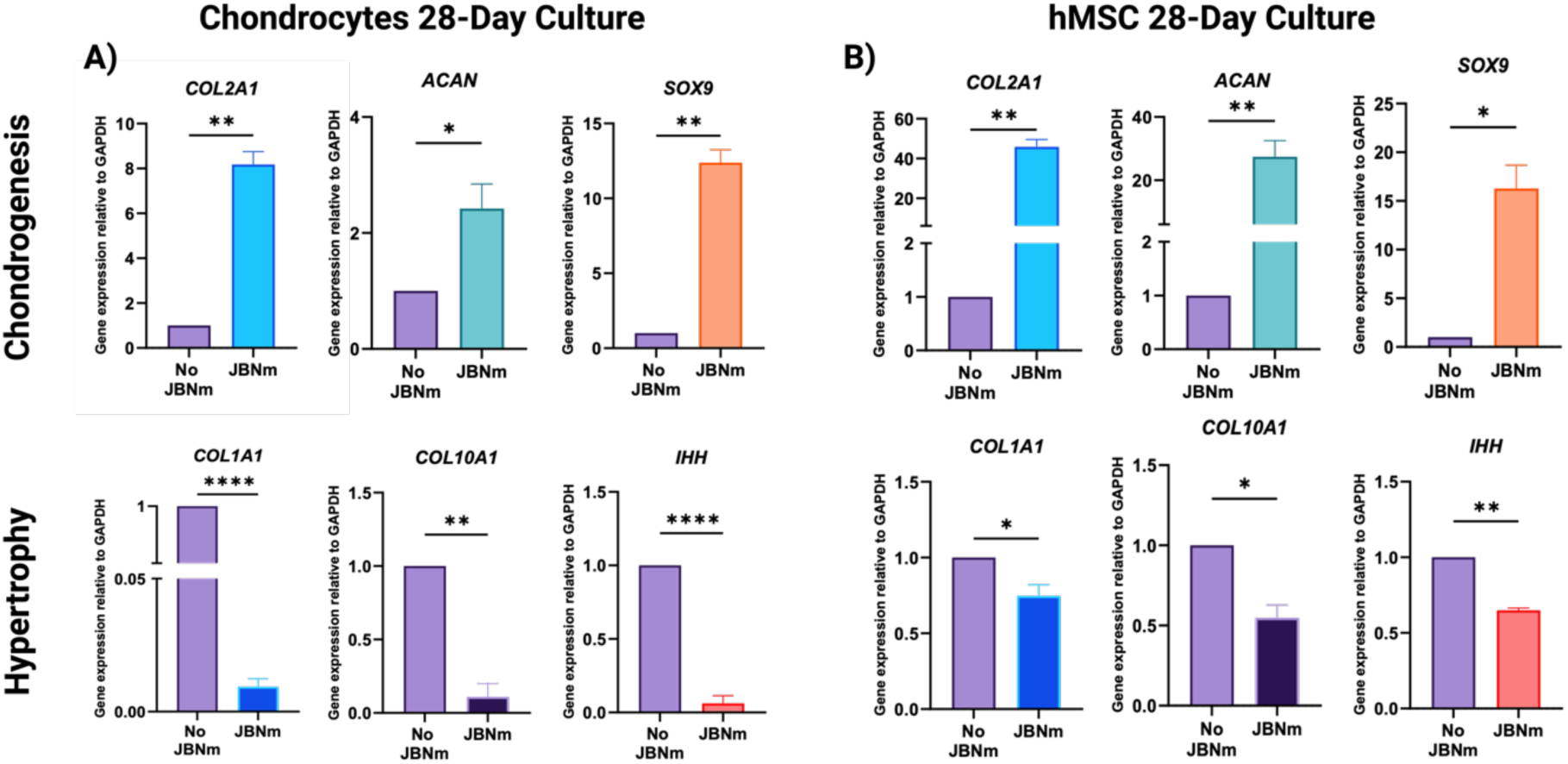
Gene expression of **(A)** cartilage cells and **(B)** hMSCs after 28 days of differentiation culture bound to JBNm versus control.

To determine the tissue reusability in tissue chips, multiple courses of IL-1α inflammatory cytokines were introduced into tissue chips (**Supplementary Figure 5**). Briefly, a cycle of 5 ng mL-1 IL-1α was administered into the tissue chips for Day 1-2 followed by no cytokine addition on Day 3-5. The cycles were repeated three times over the course of the experiment. Upon the third treatment of IL-1α, a large spike of GAG released was observed in the negative control sample when compared to the JBNm sample. We observed the recovery of tissue samples in tissue chips even after the third injection. Interestingly, we also observed the release of GAG even after the treatment was stopped after the third cycle in both groups, although the JBNm samples did decrease slightly, when compared to negative control. The turnover rate of proteoglycan is common, especially in the presence of proinflammatory cytokines, the results of this experiment show that JBNm resisted the presence of proinflammatory cytokines and can therefore mimic the underlying cartilage tissue [28]. JBNm has shown the ability to maintain homeostasis before the deterioration from biochemical-based stimulation (IL-1α) starts to affect individuals even after 15 days on consistent “treatment” of pro-inflammatory cytokines.

## Conclusion

In this study, we developed a library of biomimetic cartilage regeneration matrices using JBNts and cartilage-specific ECM proteins. Depending on the proteins encapsulated within the JBNm, we achieved cell-type binding specificity to increase stem cell preferential adhesion, proliferation, and differentiation. Furthermore, we demonstrated the impact of various ECM protein conjugates on chondrogenic and hypertrophic gene expression, identifying the best candidate for long-term cartilage regeneration while minimizing the risk of chronic de-differentiation and hypertrophy. Finally, we leveraged the mechanical characteristics of JBNm as an solid injectable scaffold for use in three-dimensional tissue-chip models for cartilage regeneration and therapeutic drug screening. JBNm provided superior chondrogenic differentiation outcomes while suppressing hypertrophic gene expression, providing efficient cartilage tissue regeneration and excellent homeostatic ECM maintenance in monolayer and tissue-on-a-chip applications. JBNm provides a promising platform for long-term cartilage maintenance and serves as an excellent platform for future cartilage-specific therapeutic drug screening.

## Methods

### JBNm fabrication and characterization

As seen in Figure 1B, a schematic diagram of JBNm fabrication composed of JBNts and Matn-1 peptide, an ECM protein commonly found in cartilage tissues. JBNts were synthesized as published previously [29]. An optimal ratio of Matn-1 and JBNts were obtained from a series of characterization experiments. The Matn-1, JBNts and resulting self-assembled Janus base Nano Matrix (JBNm) were observed and characterized by light microscopy, UV-absorbance measurement, zeta potential and transmission electron microscopy (TEM) as seen in Figure 2.

### UV-Vis absorption spectra measurement

Five groups of samples were prepared. i) For the JBNt group, 5 µL of 1 mg mL^−1^JBNts was added to 100 µL H_2_O to obtain a 47.6µg mL^−1^JBNt solution; ii) for the Matn-1 group, 50 µL of 100 µg mL^−1^Matn-1 was added to 55 µL H2O to obtain a 47.6 µg mL^−1^ Matn-1 solution; iii) for the JBNm 5:1, 100 µL of 100 µg mL^−1^ Matn-1 was mixed with 5 µL of 1 mg mL^−1^ JBNts was added to the mixture solution and pipetted several times; iv) for the JBNm 10:1, 50 µL of 100 µg mL^−1^ Matn-1 was mixed with 5 µL of 1 mg mL^−1^ JBNts and 50 µL of H_2_O was added together and pipetted several times. v) for the JBNm 20:1, 25 µL of 100 µg mL^−1^ Matn-1 was mixed with 5 µL of 1 mg mL^−1^ JBNts and 75 µL of H_2_O was added together and pipetted several times. The final concentrations of the JBNts and Matn-1 samples were 47.6 µg mL^−1^. The absorption spectrum of each group of samples was measured with a NanoDrop spectrophotometer.

### Transmission electron microscope (TEM) characterization

Four samples were prepared. i) First, 10 µL of 1 mg mL^−1^ JBNts was diluted with 40 µL distilled water to obtain a 200 µg mL^−1^ JBNt solution. For 5:1 JBNm solution, 200 µL of 100 µg mL^−1^ Matn-1 was mixed with 10 µL of 1 mg mL^−1^ JBNts to prepare the JBNm sample. For 10:1 JBNm solution, 100 µL of 100 µg mL^−1^ Matn-1 was mixed with 10 µL of 1 mg mL^−1^ JBNts to prepare the JBNm sample. Finally, for 20:1 JBNm 50 µL of 100 µg mL^−1^ Matn-1 was mixed with 10 µL of 1 mg mL^−1^ JBNts to prepare the JBNm sample. Four pieces of grids were cleaned with a Plasma Cleaner Harrick Plasma PDC-32G before negative staining was performed. The negative staining process was carried out for the specimens as follows: 3 µL JBNt solution (200 µg mL^−1^) and 3 µL of each JBNm solution were each dropped on separate grids and left for 2 min. Then, 100 µL acetate solution (0.5%) was pipetted onto the solution to rinse each grid. Excess solution was removed from the grids with filter paper and the grids were air-dried. Finally, Lab6 20-120 TEM was carried out for specimen characterization.

### Zeta-potential test

Four groups of samples were prepared. i) For the Matn-1 group, 100 µL of 100 µg mL^−1^ Matn-1 was dispersed in 700 µL H_2_O to obtain an 800 µL test solution. ii) For 5:1 JBNm, 200 µL of 100 µg mL^−1^ Matn-1 was mixed with 10 µL of 1 mg mL^−1^ JBNts and pipetted several times then dispersed in 590 µL H_2_O to obtain an 800 µL test solution. iii) 100 µL of 100 µg mL^−1^ Matn-1 was mixed with 10 µL of 1 mg mL^−1^ JBNts was added and pipetted several times then dispersed in 690 µL H_2_O to obtain an 800 µL test solution to obtain 10:1 ratio of JBNm. iv) Finally, 50 µL of 100 µg mL^−1^ Matn-1 was mixed with 10 µL of 1 mg mL^−1^ JBNts and pipetted several times to 740 µL H_2_O to obtain an 800 µL test solution 20:1 JBNm. The zeta-potential values of the two groups of samples were tested with a ZS.

### Fabrication of Tissue Chip Device

The microfluidic device contains PDMS Sylgard 184 Elastomer Kit came with Part A, an elastomer base and Part B elastomer curing agent. They are mixed 10:1 ratio (A: B) respectively for 3 minutes before putting into 4◦C to de-bubble for 2 hours. The resin was then poured onto the previously 3-D printed mold and let cure at room temperature for 72 hours. Cured tissue chips made of PDMS were then affixed onto the glass slide via uncured PDMS Sylgard 184 mixture and pressure of 1 atmosphere applied for 72 hours at room temperature seen in Figure 3. After device fabrication and assembly, the tubing and channels were sterilized by flowing 70% (v/v) ethanol through the device and drying the entire system under sterile condition for 24 hours before exposing to ultraviolet (UV) light for 30 minutes.

### Monolayer Cell culture

Concentration of JBNts is constant at 1 mg mL^−1^ and concentration of Matn-1 is 100 µg mL−1. Throughout the study, controls scaffolds refer to negative control (NC) that do not contain any embedded JBNm. Briefly, three different protein doses (10 µg mL^−1^, 1 µg mL^−1^ and 0.5 µg mL^−1^) with constant JBNts dose (10 µg mL^−1^) denoting ratio 5:1, 10:1 and 20:1, were used in this study. The JBNms were coated on the bottoms of wells of a 96 wells plate by mixing Matn-1 at 100 µg mL^−1^ and JBNt at 1 mg mL^−1^. The plate was frozen at −80 ◦C and lyophilized overnight.

### Cytotoxicity assay

Human chondrocyte cells were seeded onto two separate 96-well plates. Each well of the plates received 100 µL of cell suspension containing 3,000 cells. The two plates were incubated in a cell culture incubator for 24 h (37 ◦C, 5% CO2). Each well then received 100 µL of various concentrations of JBNts diluted in distilled water. The JBNt concentration gradient was set as 5 µg mL^−1^, 1 µg mL^−1^, 0.5 µg mL^−1^, and 0 µg mL^−1^. For each group, Similarly, three different protein doses (10 µg mL^−1^, 1 µg mL^−1^ and 0.5 µg mL^−1^) were set for the cytotoxicity assay. 6 wells were used for testing. After a 24 h incubation, each well received 10 µL of CCK-8 solution and were incubated for an additional 2 h. The absorption values of the plates were measured with multi-mode microplate readers at 450 nm.

### Adhesion test on pre-coated coverglass chambers

Three chambered coverglasses were prepared for the adhesion experiments. Each chambered coverglass included four groups of samples. For the JBNt group, 10 µL of 1 mg mL^−1^ JBNts was diluted with 190 µL distilled water to obtain a 200 µL solution. The concentration of the JBNt solution was 50 µg mL−1. For the Matn-1 group, 10 µL of 100 µg mL^−1^ Matn-1 was diluted with 190 µL distilled water to obtain a 5 µg mL^−1^ solution. For the 10:1 JBNm group, 10 µL of 10 µg mL^−1^ Matn-1 was mixed with 10 µL 1 mg mL^−1^ JBNts was added to the mixture solution and pipetted. Then, 180 µL distilled water was added to obtain a 200 µL JBNm solution. As a control group, 200 µL distilled water was used. Each sample group was added into one well of a Number 1.5 chambered coverglass. The chambered coverglasses were placed into a −80 ◦C freezer for one hour and then freeze-dried with a lyophilized instrument.

### Cell culture in tissue chips

Two groups of samples were used in this experiment, the negative control with no additives and the JBNm group at 10:1 ratio. For the 10:1 JBNm group, 20 µL of 10 mg mL^−1^ JBNts was mixed with 20 µL of 100 µg mL^−1^ Matn-1 and pipetted several times, then diluted with 260 µL cells and cell culture media, DMEM bringing it to a total of 300 µL. Three replicates were performed for each cell morphology, cell viability, cell proliferation, biochemical analysis, and gene expression for functional study of the cartilage cells.

2% agarose was obtained by mixing 200 mg agarose into 10 mL of DMEM media without serum and heated to boil to dissolve agarose completely. Cells with density of 500,000 cells/300 µL were mixed with 2% agarose (300 µL) and warmed at 40 °C to form cells/ 1% agarose mixture denoting negative control. A 300 µL mixture of cell culture with 10:1 JBNm solution and 1% agarose gel (300 µL) were injected to one tissue chip. The agarose gel with cells in both chips was then allowed cooled down to solidified at room temperature for 5 minutes under sterilized cell culture hood. Then, media was injected into the side channels allowing the cells were to form stable attachment in the chip for 1 hour in a humidified atmosphere of 5% CO_2_ in air at 37 °C. In the static group, the media exchange of the side channels was performed every other day with no disruption or application of dynamic environment for the 3D cell culture in the middle channel. In the flow group, the medium of the two chips was continuously perfused to the chip at a flow of 250 µL h^−1^ using a commercial syringe pump (Chemyx Fusion 200). The tissue chip cultures were incubated for 1,3, 5 and 7 days in a humidified atmosphere of 5% CO_2_ in air at 37 °C.

Similar methods were implemented for a long-term cell culture experiment, on hMSCs and cartilage cells. Briefly, two groups of samples were used in this experiment, the negative control with no additives and the JBNm group at 10:1 ratio. 2% agarose were combined with cells and JBNm (PBS for NC) to make 1% agarose and then injected into the middle channel of the tissue chips. Then, media were flowed into the two side channels at the rate of 250 µL h^−1^ using a commercial syringe pump and cultured for 28 days in a humidified atmosphere of 5% CO_2_ in air at 37 °C.

### Tissue chip validation with JBNm

Two sample groups in triplicates were used in this experiment with 8×106 cells mL^−1^ in a humidified atmosphere of 5% CO_2_ in air at 37 °C. The first group is the negative control with no additives and t the JBNm group at the 10:1 ratio with the addition of Interleukin-1-alpha (IL-1α) at 5 ng mL−1. IL-1α was added into one medium channel, while a standard DMEM media was added into another medium channel into a chip with 1% agarose containing injectable JBNm scaffold and cells then ran at 250 h^−1^ for 12-, 24- and 48-hours. Media sample was collected from the reservoir after each time point. A negative control of the chip was also assessed without JBNm.

Similarly, in another group, 5ng mL^−1^ IL-1α was added together with 500 mL^−1^ Interleukin-1-Receptor antagonists (IL-Ra) to determine the applicability of the tissue chip with the presence of JBNm. Samples were perfused with 250 µL h^−1^ using standard media in one channel and IL-1α and IL-1Ra media in another channel for 12-, 24- and 48-hours continuously. Supernatants are collected after each time point. A negative control of the chip was also assessed without JBNm.

### Cell Viability Analysis in Tissue Chips

Cell viability on tissue chips in static condition compared to flow condition at 250 µL h^−1^ were quantified (Figure 5). After each time point at 1,3, 5 and 7 days, the media from side channels were aspirated, and washed with PBS. The PDMS chip was peeled off carefully under the sterilized condition in the fume hood and transferred to an 8-well chambered coverglass slide for further processing. After samples were washed with PBS, Live Dead Cell Viability Assay Kit for 3D and 2D cell culture CBA415 was used according to the manufacturer protocol. The viability of cells in tissue chips were assessed and analyzed with A Nikon A1R spectral confocal microscope and finally quantified with ImageJ.

### Cell Morphology Analysis in Tissue Chips

After each time point, the media from side channels were aspirated, and washed with PBS. Then, the chips were peeled away from the attached glass slide to access the cells embedded in agarose. Then, the cell embedded agarose was transferred to a chambered coverglass before fixing with 4% paraformaldehyde for 30 minutes. The fixative solution was removed, the cells were rinsed twice with PBS, and the cells were incubated with 100 µL 0.1% Triton-X for 10 minutes. After two rinses with PBS, 100 µL of 0.165 µM rhodamine-phalloidin was added to each well for 30 minutes. Next, 0.1 µg mL^−1^ DAPI was used to stain the cell nuclei. After a 5 minutes incubation, the DAPI was pipetted out. Finally, the cells were rinsed twice with PBS. A Nikon A1R spectral confocal microscope was used to observe the morphology and obtain fluorescent images of cells. Analyses of the number and morphology of cells were performed with Cell Profiler, and ImageJ.

### Cell proliferation and Biochemical Assay

Similarly, after each time point, the chips were peeled away from the glass slide and transferred to microcentrifuge tubes. The cells embedded agarose were washed with PBS and incubated with trypsin for 30 minutes at room temperature to detach cells from the agarose. After centrifuge, trypsin was aspirated, and cells were re-suspended in PBS containing 50 µg mL^-1^ proteinase K (Sigma) and digested for 16 hours at 56 °C. The DNA content reflects cell proliferation was tested by using Hoechst 33258 assay. In each sample, absorbance (460 nm) of total intracellular DNA content, as indicated by the Hoechst 33258 dye, was measured using a fluorescence microplate Eppendorf, Hamburg, Germany. The production of glycosaminoglycans (GAGs) was determined by 1,9-dimethylmethylene blue (DMMB) method. Absorbance (530 nm) of total GAG secretion was measured by the microplate reader and converted into concentration using chondroitin sulfate standard curve. GAG content in each cell was normalized to the total DNA content of all chondrocytes that showed their activity in the agarose gel culture in the chip.

### Tissue chip drug screening and reusability

The setup for this experiment is similar to the setup with cell culture in tissue chip methods described above. For tissue chip drug screening, JBNm and cartilage cells are cultured in the five tissue chips for 24 hours, followed by addition of 5 ng mL^-1^ of IL-1α in media for another 24 hours before addition of interleukin-1 receptor antagonist (IL-1Ra) of different concentrations, starting with 5 ng mL^-1^, 20 ng mL^-1^, 40 ng mL^-1^, 100 ng mL^-1^ and 200 ng mL^-1^ for the next 3 days for n=1. Repeat of n=2 and n=3 was conducted as well with similar experimental condition. Similarly, resveratrol with different concentrations at 0.1 µg mL^-1^, 0.5 µg mL^-1^, 2 µg mL^-1^, 10 µg mL^-1^ and 20 µg mL^-1^ with 3 repeats. JBNm in tissue chips, as well as NC without JBNm were also conducted. Then, supernatants were collected every 24 hours and GAG degradation was determined by DMMB method at absorbance of 530 nm read in microplatereader. Next, IC_50_ was also determined for both IL-1Ra and Resveratrol. Finally, qRT-PCR was performed from all samples with methods described below.

For tissue chip reusability, after 24 hours of cartilage cells maturing in six tissue chips, JBNm samples (three tissue chips) and negative samples (three tissue chips) were treated with 5 ng mL^-1^ of IL-1α in media for 48 hours in the superficial zone and the pro-inflammatory cytokine was taken off for 72 hours, denoting the first cycle. A total of three cycles were conducted and supernatant were collected at selected timepoint to determine the release of GAG denoting the degradation of GAG in both superficial and deep zone of the tissue chip and determined by DMMB method at absorbance of 530 nm read in microplate reader.

### RNA extraction and qRT-PCR

Gene expressions of ACAN, COL II, and SOX9 were detected by qRT-PCR using primers sequence found in table 1. Total RNA extraction was performed using the RNA isolation kit according to the manufacturer’s instructions. A reverse transcription kit was used for reverse transcription of RNA, as was carried out at 25 °C for 5 min, 42 °C for 60 min and then 72 °C for 5 min. qRT-PCR was performed using Fast Start Universal SYBR Green Master under the condition of 10 min at 95 °C, 15 s at 95 °C and 1 min at 0 °C. The dissociation curve of each primer pair was analyzed to confirm primer specificity. Fold increase in PCR products was analyzed using a 2−ΔΔCt method. Data was normalized to GAPDH messenger RNA levels. All experiments were performed three times for each gene.

## Acknowledgements

We would like to thank NIH 7R01AR072027, NIH R01GM155969-01, NIH 1R21AR079153-01A1, NSF 2025362, NSF 2234570 and the University of Connecticut for funding.

## References

[1] F. Causa, P. A. Netti, and L. Ambrosio, “A multi-functional scaffold for tissue regeneration: The need to engineer a tissue analogue,” Biomaterials, vol. 28, no. 34, pp. 5093–5099, 2007/12/01/ 2007, doi: 10.1016/j.biomaterials.2007.07.030.

[2] Y. Gao et al., “The ECM-Cell Interaction of Cartilage Extracellular Matrix on Chondrocytes,” BioMed Research International, vol. 2014, no. 1, p. 648459, 2014, doi: 10.1155/2014/648459.

[3] D. C. Ding, W. C. Shyu, and S. Z. Lin, “Mesenchymal stem cells,” (in eng), Cell Transplant, vol. 20, no. 1, pp. 5–14, 2011, doi: 10.3727/096368910x.

[4] A. Aldahmash, W. Zaher, M. Al-Nbaheen, and M. Kassem, “Human stromal (mesenchymal) stem cells: basic biology and current clinical use for tissue regeneration,” (in eng), Ann Saudi Med, vol. 32, no. 1, pp. 68–77, Jan-Feb 2012, doi: 10.5144/0256-4947.2012.68.

[5] D. F. Duarte Campos, W. Drescher, B. Rath, M. Tingart, and H. Fischer, “Supporting Biomaterials for Articular Cartilage Repair,” CARTILAGE, vol. 3, no. 3, pp. 205–221, 2012, doi: 10.1177/1947603512444722.

[6] I.-C. Liao, F. T. Moutos, B. T. Estes, X. Zhao, and F. Guilak, “Composite Three-Dimensional Woven Scaffolds with Interpenetrating Network Hydrogels to Create Functional Synthetic Articular Cartilage,” Advanced Functional Materials, vol. 23, no. 47, pp. 5833–5839, 2013, doi: 10.1002/adfm.201300483.

[7] Z. Mahdieh et al., “Granular Matrigel: restructuring a trusted extracellular matrix material for improved permeability,” Biomedical Materials, vol. 17, no. 4, p. 045020, 2022/06/08 2022, doi: 10.1088/1748-605X/ac7306.

[8] T. L. Vincent, O. McClurg, and L. Troeberg, “The Extracellular Matrix of Articular Cartilage Controls the Bioavailability of Pericellular Matrix-Bound Growth Factors to Drive Tissue Homeostasis and Repair,” International Journal of Molecular Sciences, vol. 23, no. 11, p. 6003, 2022.

[9] M. Riedl et al., “Attenuation of Hypertrophy in Human MSCs via Treatment with a Retinoic Acid Receptor Inverse Agonist,” (in eng), Int J Mol Sci, vol. 21, no. 4, Feb 20 2020, doi: 10.3390/ijms21041444.

[10] N. Giuliani et al., “New insights into osteogenic and chondrogenic differentiation of human bone marrow mesenchymal stem cells and their potential clinical applications for bone regeneration in pediatric orthopaedics,” (in eng), Stem Cells Int, vol. 2013, p. 312501, 2013, doi: 10.1155/2013/312501.

[11] M. D’Angelo, P. C. Billings, M. Pacifici, P. S. Leboy, and T. Kirsch, “Authentic matrix vesicles contain active metalloproteases (MMP): a role for matrix vesicle-associated MMP-13 in activation of transforming growth factor-β,” Journal of Biological Chemistry, vol. 276, no. 14, pp. 11347–11353, 2001.

[12] M. D’Angelo et al., “MMP-13 is induced during chondrocyte hypertrophy,” Journal of cellular biochemistry, vol. 77, no. 4, pp. 678–693, 2000.

[13] R. A. Somoza, J. F. Welter, D. Correa, and A. I. Caplan, “Chondrogenic differentiation of mesenchymal stem cells: challenges and unfulfilled expectations,” (in eng), Tissue Eng Part B Rev, vol. 20, no. 6, pp. 596–608, Dec 2014, doi: 10.1089/ten.TEB.2013.0771.

[14] H. Chen et al., “Molecular Mechanisms of Chondrocyte Proliferation and Differentiation,” (in eng), Front Cell Dev Biol, vol. 9, p. 664168, 2021, doi: 10.3389/fcell.2021.664168.

[15] N. Ma, X. Teng, Q. Zheng, and P. Chen, “The regulatory mechanism of p38/MAPK in the chondrogenic differentiation from bone marrow mesenchymal stem cells,” Journal of Orthopaedic Surgery and Research, vol. 14, no. 1, p. 434, 2019/12/12 2019, doi: 10.1186/s13018-019-1505-2.

[16] A. Mansour, M. Romani, A. B. Acharya, B. Rahman, E. Verron, and Z. Badran, “Drug Delivery Systems in Regenerative Medicine: An Updated Review,” (in eng), Pharmaceutics, vol. 15, no. 2, Feb 18 2023, doi: 10.3390/pharmaceutics15020695.

[17] A. Yau, Y. Chen, and M. Landolina, “Fabrication and Characterization of Layer-By-Layer Janus Base Nano-Matrix to Promote Cartilage Regeneration,” JoVE, no. 185, p. e63984, 2022/07/06 2022, doi: doi:10.3791/63984.

[18] F. Deák, R. Wagener, I. Kiss, and M. Paulsson, “The matrilins: a novel family of oligomeric extracellular matrix proteins,” Matrix Biology, vol. 18, no. 1, pp. 55–64, 1999/02/01/ 1999, doi: 10.1016/S0945-053X(98)00006-7.

[19] K. L. Posey, F. Coustry, and J. T. Hecht, “Cartilage oligomeric matrix protein: COMPopathies and beyond,” Matrix Biology, vol. 71-72, pp. 161–173, 2018/10/01/ 2018, doi: 10.1016/j.matbio.2018.02.023.

[20] J. A. Martin and J. A. Buckwalter, “Effects of fibronectin on articular cartilage chondrocyte proteoglycan synthesis and response to insulin-like growth factor-I,” Journal of Orthopaedic Research, vol. 16, no. 6, pp. 752–757, 1998, doi: 10.1002/jor.1100160618.

[21] E. Leclerc, Y. Sakai, and T. Fujii, “Cell Culture in 3-Dimensional Microfluidic Structure of PDMS (polydimethylsiloxane),” Biomedical Microdevices, vol. 5, no. 2, pp. 109–114, 2003/06/01 2003, doi: 10.1023/A:1024583026925.

[22] H. Harjunpää, M. Llort Asens, C. Guenther, and S. C. Fagerholm, “Cell Adhesion Molecules and Their Roles and Regulation in the Immune and Tumor Microenvironment,” (in English), *Frontiers in Immunology*, Review vol. 10, 2019-May-22 2019, doi: 10.3389/fimmu.2019.01078.

[23] C. Jensen and Y. Teng, “Is It Time to Start Transitioning From 2D to 3D Cell Culture?,” (in English), *Frontiers in Molecular Biosciences*, Review vol. 7, 2020-March-06 2020, doi: 10.3389/fmolb.2020.00033.

[24] R. J. Smith, N. A. Rohloff, L. M. Sam, J. M. Justen, M. R. Deibel, and J. C. Cornette, “Recombinant human interleukin-1α and recombinant human interleukin-1β stimulate cartilage matrix degradation and inhibit glycosaminoglycan synthesis,” Inflammation, vol. 13, no. 4, pp. 367–382, 1989/08/01 1989, doi: 10.1007/BF00914921.

[25] L. Zhou, W. Zhang, J. Lee, L. Kuhn, and Y. Chen, “Controlled Self-Assembly of DNA-Mimicking Nanotubes to Form a Layer-by-Layer Scaffold for Homeostatic Tissue Constructs,” ACS Applied Materials & Interfaces, vol. 13, no. 43, pp. 51321–51332, 2021/11/03 2021, doi: 10.1021/acsami.1c13345.

[26] P. Hoff et al., “Osteoarthritis synovial fluid activates pro-inflammatory cytokines in primary human chondrocytes,” International Orthopaedics, vol. 37, no. 1, pp. 145–151, 2013/01/01 2013, doi: 10.1007/s00264-012-1724-1.

[27] M. D. Turner, B. Nedjai, T. Hurst, and D. J. Pennington, “Cytokines and chemokines: At the crossroads of cell signalling and inflammatory disease,” Biochimica et Biophysica Acta (BBA) - Molecular Cell Research, vol. 1843, no. 11, pp. 2563–2582, 2014/11/01/ 2014, doi: 10.1016/j.bbamcr.2014.05.014.

[28] M. A. Karsdal, S. H. Madsen, C. Christiansen, K. Henriksen, A. J. Fosang, and B. C. Sondergaard, “Cartilage degradation is fully reversible in the presence of aggrecanase but not matrix metalloproteinase activity,” Arthritis Research & Therapy, vol. 10, no. 3, p. R63, 2008/05/30 2008, doi: 10.1186/ar2434.

[29] H. Fenniri et al., “Helical rosette nanotubes: design, self-assembly, and characterization,” (in eng), J Am Chem Soc, vol. 123, no. 16, pp. 3854–5, Apr 25 2001, doi: 10.1021/ja005886l.

