## Supplemental Data for "Injectable Janus Base Nanomatrix (JBNm) in Maintaining Long-Term Homeostasis of Regenerated Cartilage for Tissue Chip Applications"

### Slide 1
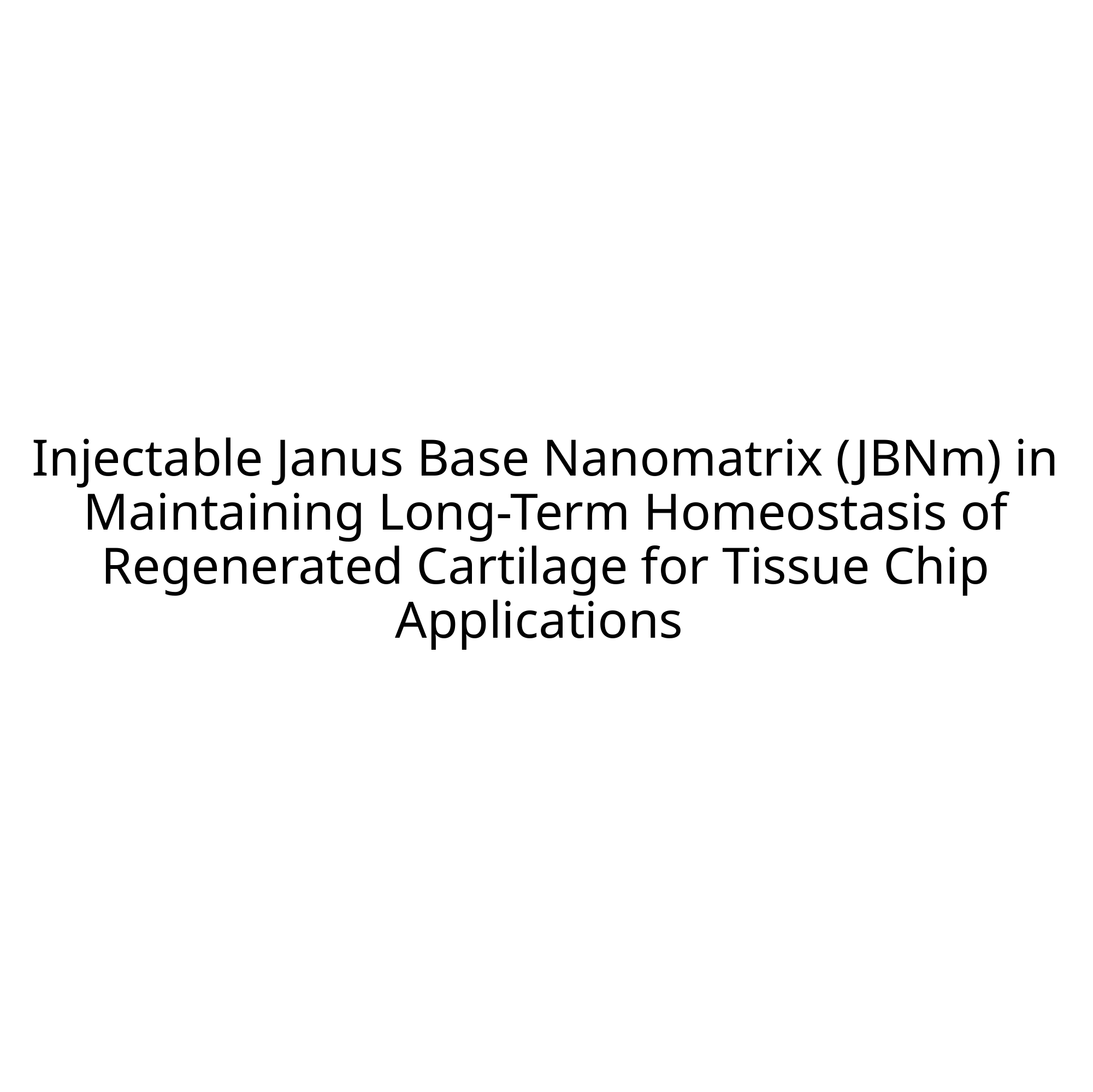

#### Injectable Janus Base Nanomatrix (JBNm) in Maintaining Long-Term Homeostasis of Regenerated Cartilage for Tissue Chip Applications

### Slide 2
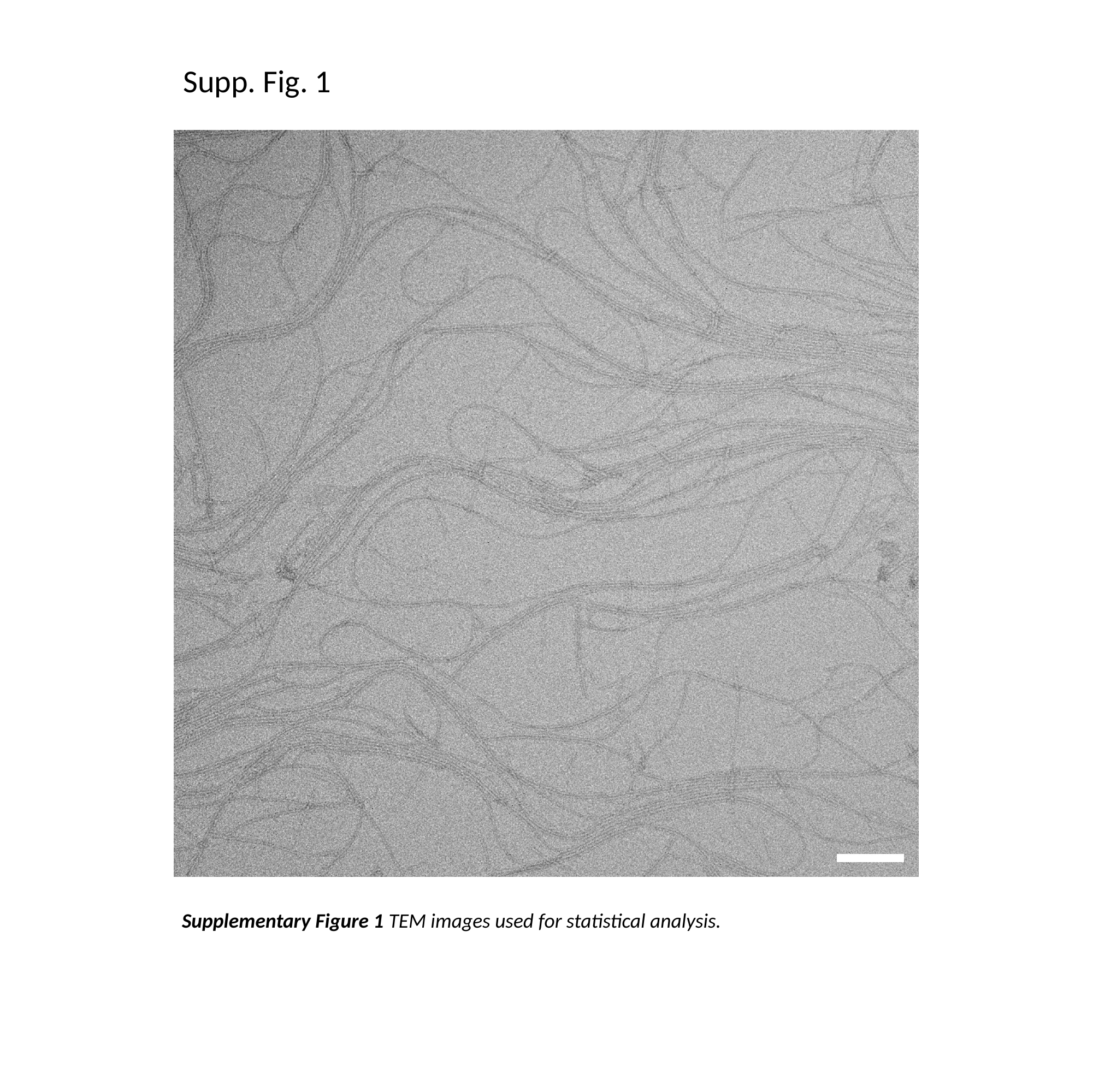

Supp. Fig. 1
Supplementary Figure 1 TEM images used for statistical analysis.

### Slide 3
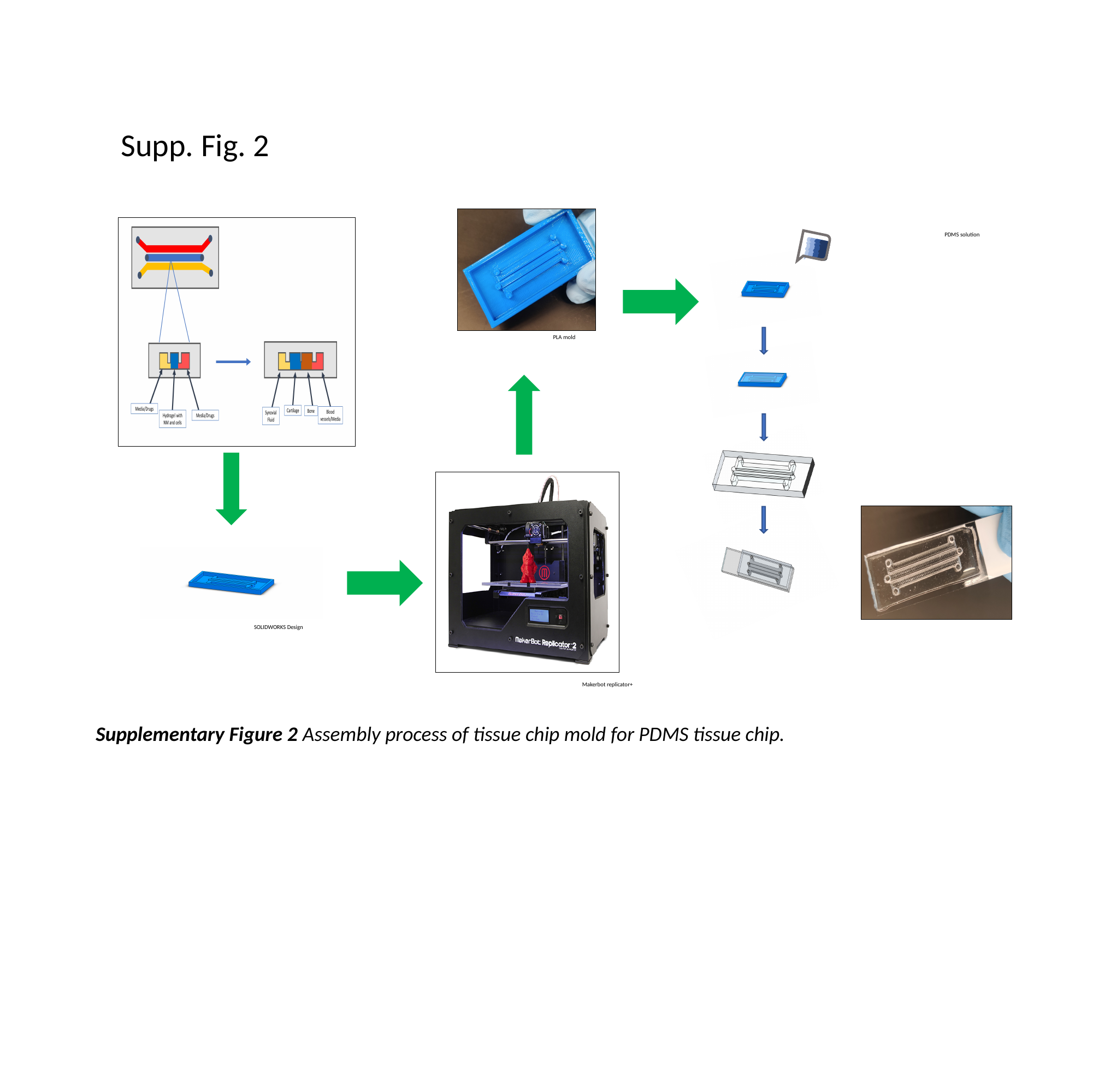

Supp. Fig. 2
PDMS solution
PLA mold
SOLIDWORKS Design
Makerbot replicator+
Supplementary Figure 2 Assembly process of tissue chip mold for PDMS tissue chip.

### Slide 4
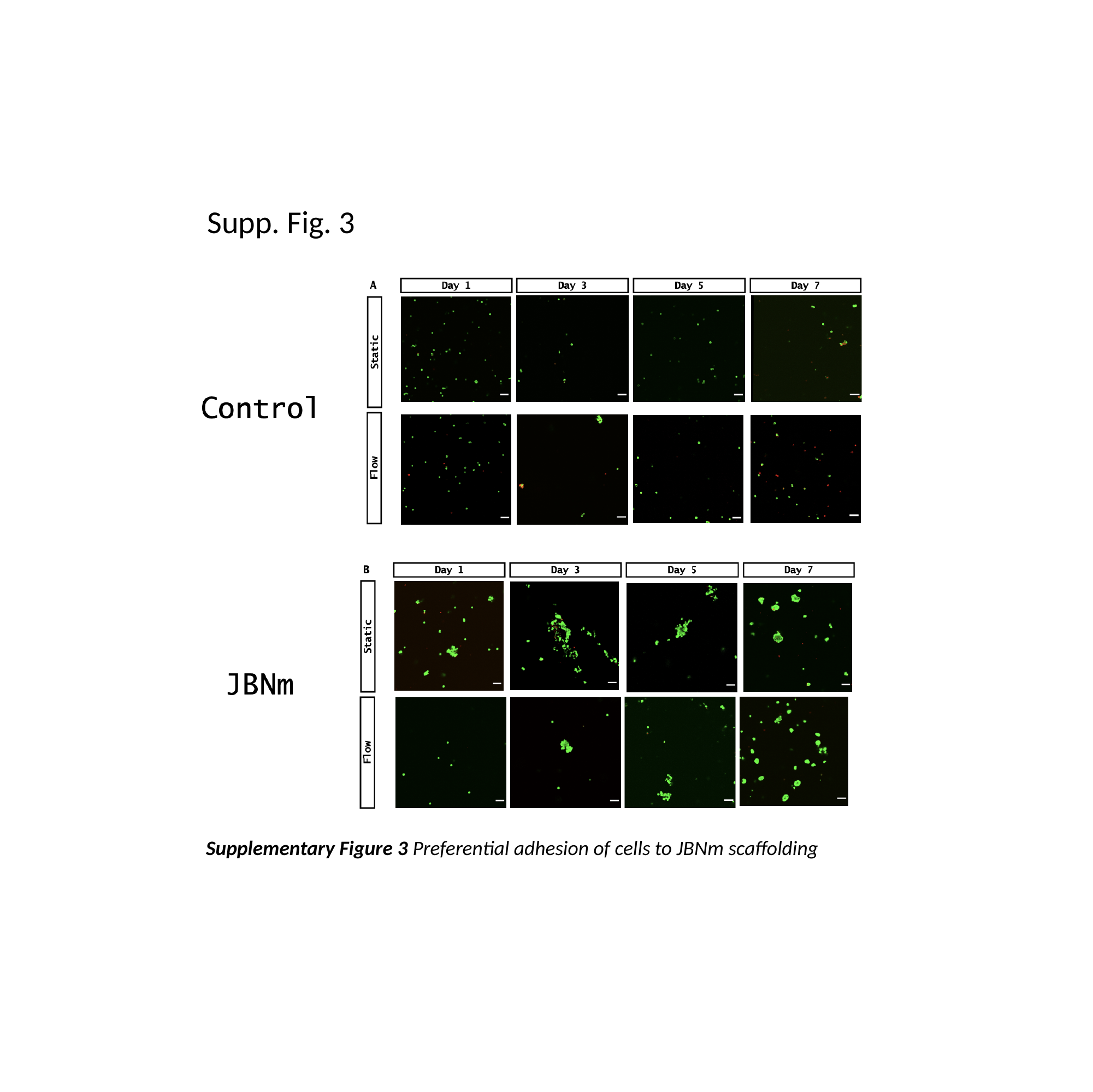

Supp. Fig. 3
Supplementary Figure 3 Preferential adhesion of cells to JBNm scaffolding

### Slide 5
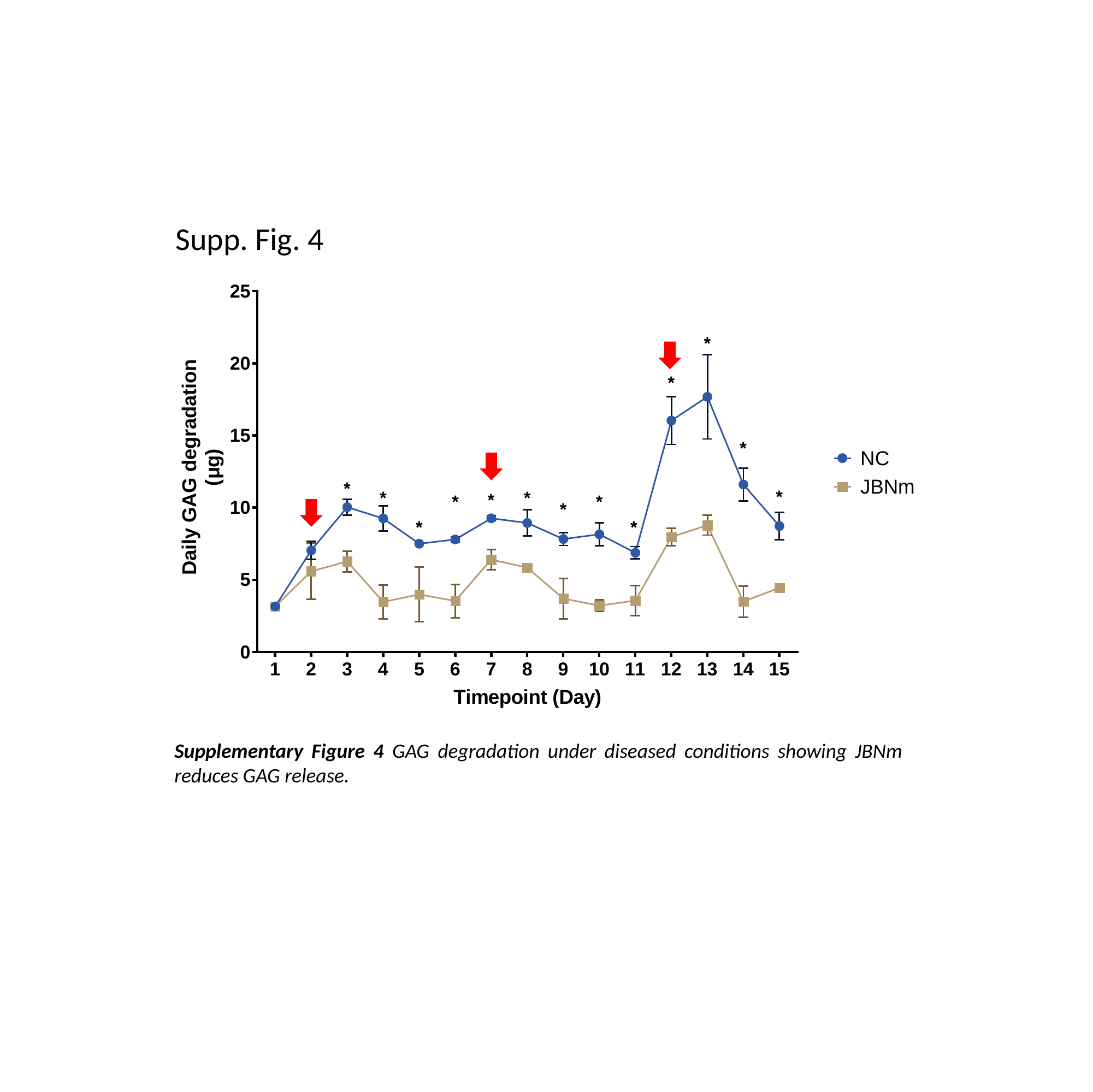

Supp. Fig. 4
Supplementary Figure 4 GAG degradation under diseased conditions showing JBNm reduces GAG release.

### Slide 6
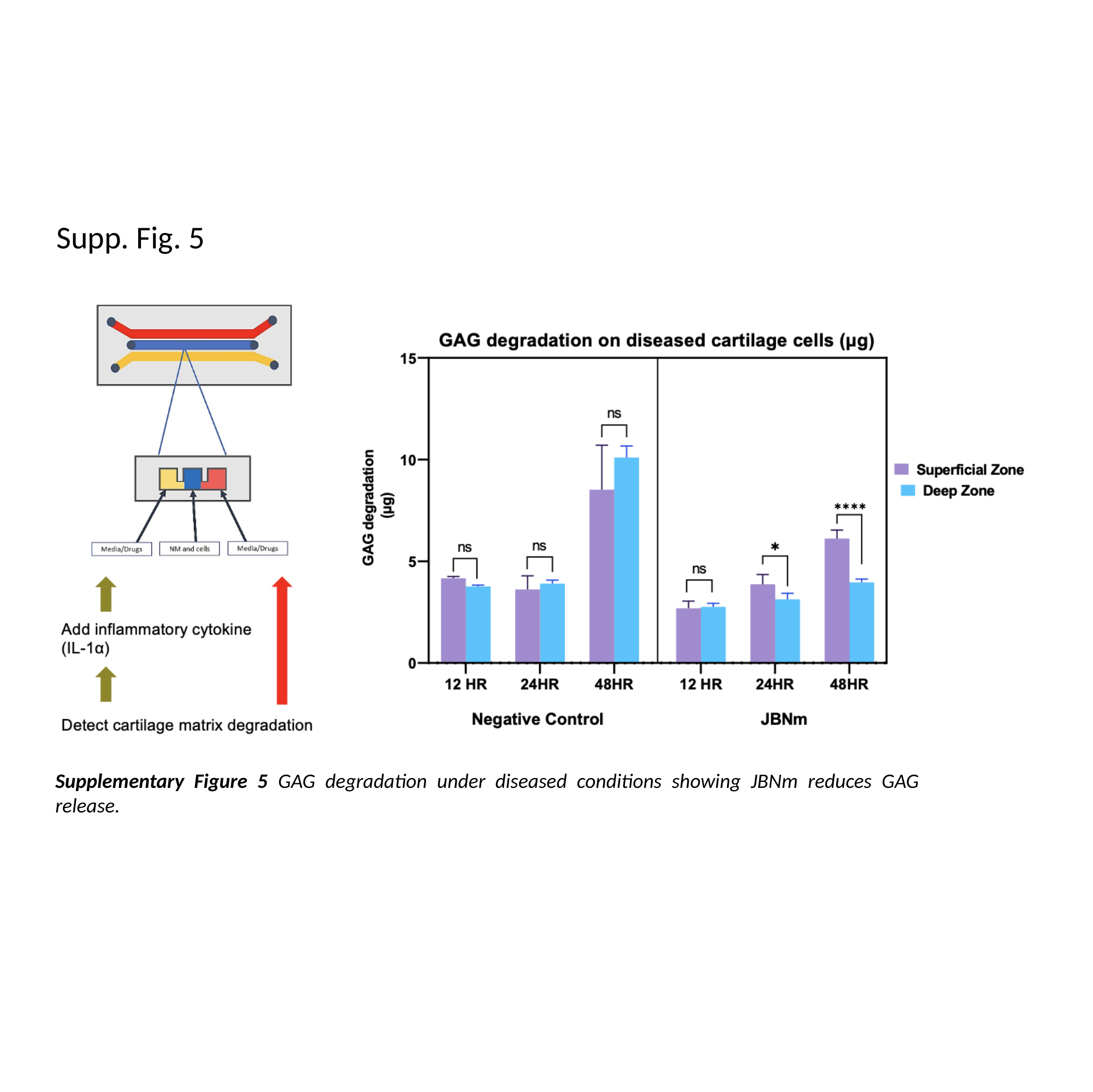

Supp. Fig. 5
Supplementary Figure 5 GAG degradation under diseased conditions showing JBNm reduces GAG release.

### Slide 7
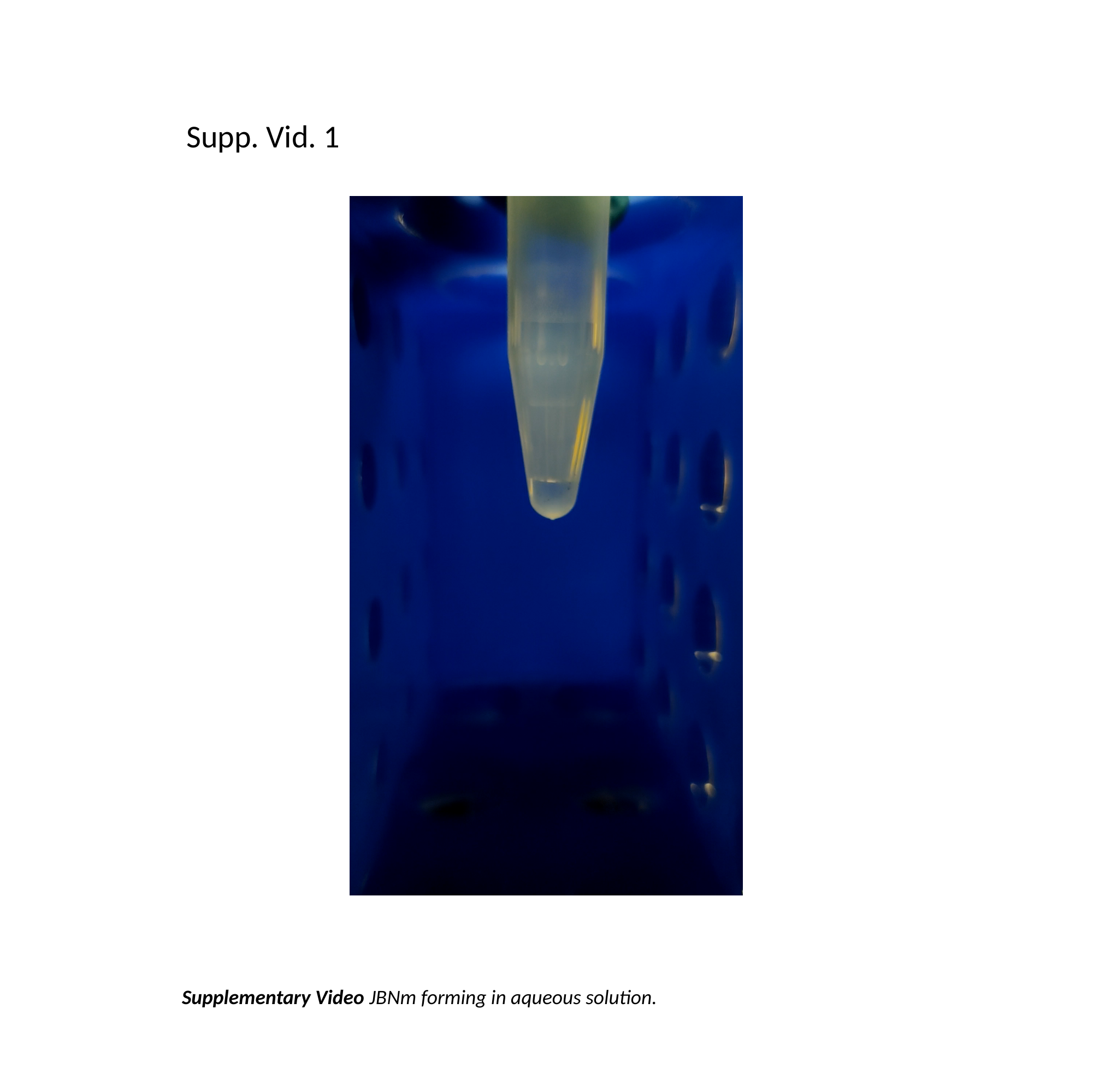

Supp. Vid. 1
Supplementary Video JBNm forming in aqueous solution.
